# The effect of a dominant kinase-dead *Csf1r* mutation associated with adult-onset leukoencephalopathy on brain development and neuropathology

**DOI:** 10.1101/2024.06.12.598773

**Authors:** Jennifer Stables, Reiss Pal, Barry M. Bradford, Dylan Carter-Cusack, Isis Taylor, Clare Pridans, Nemat Khan, Trent Woodruff, Katharine M. Irvine, Kim M. Summers, Neil A. Mabbott, David A. Hume

## Abstract

Amino acid substitutions in the kinase domain of the human *CSF1R* protein are associated with autosomal dominant adult-onset leukoencephalopathy with axonal spheroids and pigmented glia (ALSP). To model the human disease, we created a disease-associated mutation (Glu631Lys; E631K) in the mouse *Csf1r* locus. Previous analysis demonstrated that heterozygous mutation (*Csf1r*^E631K/+^) had a dominant inhibitory effect on CSF1R signaling *in vitro* and *in vivo* but did not recapitulate the pathology of the human disease. We speculated that leukoencephalopathy in humans requires an environmental trigger and/or epistatic interaction with common neurodegenerative disease-associated alleles. Here we examine the impact of heterozygous *Csf1r* mutation on microglial phenotype, normal postnatal brain development, age-related changes in gene expression and on two distinct pathologies in which microgliosis is a prominent feature, prion disease and experimental autoimmune encephalitis (EAE). The heterozygous *Csf1r*^E631K/+^ mutation reduced microglial abundance and the expression of microglial-associated transcripts relative to wild-type controls at 12 weeks and 43 weeks of age but had no selective effect on homeostatic markers such as *P2ry12*. An epistatic interaction was demonstrated between *Csf1r*^E631K/+^ and *Cxc3r1*^EGFP/+^ genotypes leading to dysregulated microglial and neuronal gene expression in both hippocampus and striatum. Heterozygous *Csf1r*^E631K^ mutation reduced the microgliosis associated with both diseases. There was no significant impact on disease severity or progression in prion disease. In EAE, induced expression of inflammation-associated transcripts in the hippocampus and striatum was suppressed in parallel with microglia-specific transcripts, but spinal cord demyelination was exacerbated. The results support a dominant-negative model of CSF1R-associated leukoencephalopathy and likely contributions of an environmental trigger and/or genetic background to neuropathology.

## Introduction

Signals from the colony stimulating factor 1 receptor (CSF1R) are required for survival, proliferation, and differentiation of mononuclear phagocyte populations throughout the body including microglia in the brain (Chitu and Stanley, 2017; Hume et al., 2020; Keshvari et al., 2021; Patkar et al., 2021). CSF1R has two ligands, colony stimulating factor 1 (CSF1) and interleukin 34 (IL34) (Lelios et al., 2020). Upon ligand binding, CSF1R dimerization and autophosphorylation generates phosphotyrosine docking sites for multiple downstream effector pathways (Chitu and Stanley, 2017; Stanley and Chitu, 2014). Biallelic recessive loss-of-function mutations in mouse, rat and human *CSF1R* genes are causally linked to osteopetrosis and postnatal developmental abnormalities (reviewed in (Chitu et al., 2021; Hume et al., 2020). In 2011, Rademakers *et al* (Rademakers et al., 2011) identified amino acid substitutions in the tyrosine kinase domain of CSF1R in patients with autosomal dominant adult-onset leukoencephalopathy with axonal spheroids and pigmented glia (ALSP), now also called CSF1R-related leukoencephalopathy (CRL) (Chitu et al., 2021). More than 140 different disease-associated *CSF1R* coding mutations have been identified (Chitu et al., 2021; Guo and Ikegawa, 2021; Konno et al., 2018; Konno et al., 2017; Oosterhof et al., 2019).

We recently published the characterisation of a transgenic mouse line with a disease-associated *Csf1r* mutation (substitution of lysine for glutamine at position amino acid position 631, abbreviated to E631K, equivalent to E633K in humans). Like human patients with heterozygous *CSF1R* mutations (Tada et al., 2016), heterozygous mutant mice (*Csf1r*^E631K/+^) had reduced density of microglia with altered morphology. They were less responsive to CSF1-induced bone marrow cell proliferation *in vitro* and to treatment with a CSF1-Fc fusion protein *in vivo* (Stables et al., 2022). These findings supported a dominant-negative model for ALSP in which mutant receptors are expressed on the cell surface, bind and internalise the ligand, but compromise signalling from the receptor produced by the wild-type (WT) allele (Pridans et al., 2013). Further support for a dominant-negative model was provided by analysis of a kinase-dead *Csf1r* mutation in zebrafish and brain tissue from affected individuals (Berdowski et al., 2022). In patients, kinase-dead CSF1R homodimers and heterodimers may also compete for ligands with the functional receptor dimers (Hume et al., 2020). Similar dominant kinase-dead mutations have been reported in the closely-related *Kit* gene in mice and in human piebaldism (Oiso et al., 2013; Reith et al., 1990). The alternative disease model is that the loss of 50% of CSF1R expression or function (haploinsufficiency) is sufficient to give rise to neuropathology (Chitu et al., 2021). Heterozygous *Csf1r* mutation on a C57Bl/6J mouse genetic background gave rise to microgliosis and evidence of neuropathology with variable penetrance that depended in part upon diet (Chitu et al., 2020; Chitu et al., 2015; Chitu et al., 2021). However, although there are focal concentrations of macrophages in lesions in end-stage ALSP, the predominant feature is depletion of microglia rather than microgliosis (Berdowski et al., 2022; Oosterhof et al., 2018; Tada et al., 2016).

Microglia have been ascribed numerous roles in brain development, maturation, synaptic plasticity and in neurodegeneration (Paolicelli et al., 2022). Against that background, normal brain development in the absence of microglia in mice with a homozygous *Csf1r* enhancer mutation (*Csf1r*^ΔFIRE/ΔFIRE^) was surprising (Rojo et al., 2019). Subsequent analysis of these mice suggested that the lack of microglia led to an age-dependent loss of myelin integrity with similarities to ALSP supporting the view that microglial deficiency rather than microgliosis underlies neuropathology in patients (McNamara et al., 2023). The trophic functions of microglia were also evident in accelerated pathology observed in models of prion disease and Alzheimer’s dementia in their absence (Bradford et al., 2022; Kiani Shabestari et al., 2022). Microglia-deficient *Csf1r*^ΔFIRE/ΔFIRE^ mice show evidence of age-dependent pathology, including demyelination (McNamara et al., 2023) and vascular calcification in the thalamus that was reversed/prevented by restoration of microglia {Chadarevian, 2024 #409; Munro et al., 2024). *Csf1r^+/-^* mice were reported to exhibit neuropathologies including ventricular enlargement, thinning of the corpus callosum and axonal spheroids in common with ALSP patients (Chitu et al., 2015). Our previous analysis of the *Csf1r*^E631K/+^ mice including MRI, provided no evidence of any comparable brain pathology even up to 15 months of age (Stables et al., 2022).

Chitu *et al*. (Chitu et al., 2015) reported that by 11 weeks of age, *Csf1r*^+/-^ mice on the C57BL/6J background developed a mild microgliosis (20-30% increase) throughout the brain detected by localisation of IBA1 antigen (encoded by *Aif1*). By contrast, *Csf1r*^E631K/+^ mice showed a consistent 30-40% reduction in IBA1^+^ microglia across all brain regions (Stables et al., 2022). Arreola *et al*. (Arreola et al., 2021) reported the loss of presynaptic surrogates and perineuronal nets in *Csf1r*^+/-^ mice that was reversed by microglial elimination. Aside from the reduction in density, microglia in *Csf1r^E631K/+^* mice showed reduced dendritic arborisation (Stables et al., 2022). Altered morphology is considered an indication of reactive microglia, a feature of human ALSP (Kempthorne et al., 2020; Tada et al., 2016). In a model of Cre-induced heterozygous mutation of *Csf1r* (Arreola et al., 2021), microglial dyshomeostasis was evident from reduced expression of microglia-enriched markers such as P2RY12. By contrast, comparative immunolocalization of P2RY12 and TMEM119 in *Csf1r*^+/+^ and *Csf1r*^E631K/+^ cortex revealed no difference in the staining intensity and punctate distribution of these markers (Stables et al., 2022)

An important tool in the study of microglial biology has been the development of orally-available brain-permeant CSF1R kinase inhibitors (Green et al., 2020; Liu et al., 2021; Montilla et al., 2023). Whilst studies of these drugs can be of interest from a purely pharmacological viewpoint, there are several caveats to interpretation. Pharmaceutical elimination of microglia produces dead cells which must be cleared, and off-target effects on related kinases and impacts on the peripheral macrophage populations which also depend upon CSF1R signals cannot be excluded. Aside from the direct relevance to human disease, the heterozygous *Csf1r*^E631K/+^ mouse provides a unique alternative model in which to assess the quantitative importance of CSF1R signalling in brain homeostasis and pathology. Here we explore the impact of the dominant *Csf1r* mutation on microglial homeostasis, brain development and models of neuropathology.

## Results

### Phenotypic analysis of Csf1r^E631K/+^ mice

Microglia are so abundant in brain that the expression signature and impact of *Csf1r* loss of function can be readily detected in total tissue mRNA (Patkar et al., 2021; Rojo et al., 2019). Such analysis avoids artefacts associated with microglial isolation, including incomplete and unrepresentative recovery, activation during isolation and contamination with mRNA from unrelated cells (reviewed in (Hume et al., 2023)). To assess the impact of the heterozygous *Csf1r*^E631K^ genotype we profiled expression by RNA-seq in hippocampus and striatum at 12 weeks and 43 weeks of age. In view of the male bias reported for the *Csf1r*^+/-^ model (Chitu et al., 2015) and reported transcriptomic differences in microglia from male and female mice (Guneykaya et al., 2018; Hanamsagar et al., 2018; Villa et al., 2018), we profiled equal numbers of males and females. The complete dataset is provided in **Table S1A.** With the exception of the male-specific expression of Y chromosome-specific transcripts (*Ddx3y, Eif2s3y, Kdm5d, Uty*) the male or female-enriched expression of specific transcripts reported previously (Guneykaya et al., 2018; Hanamsagar et al., 2018; Villa et al., 2018) was not replicated in our data in hippocampus or striatum at either age. To identify co-regulated transcripts, the dataset was clustered using the network analysis software, Graphia (Freeman et al., 2022); Clusters and their patterns of shared expression (average expression of all transcripts within the cluster) are provided in **Table S1B.** The largest clusters are age-specific and/or brain region-specific and unrelated to genotype. **Cluster 1,** around 4000 transcripts, groups based upon selective down-regulation with age in both brain regions. Striatum-enriched transcripts in **Cluster 4** were also selectively down-regulated with age. However, **Cluster 13,** containing *Csf1r,* was the only cluster clearly associated with *Csf1r* genotype in both brain regions and at both ages. **Figure 1A** shows the average expression of the 36 transcripts in this cluster. They are a subset of the CSF1R-dependent transcripts identified by profiling of the brains of microglia-deficient mice (Rojo et al., 2019), in mice treated with a CSF1R kinase inhibitor (Arreola et al., 2021; Elmore et al., 2014) and in brain tissue from ALSP patients (Berdowski et al., 2022). This pattern is entirely consistent with the reduced density of IBA1^+^ microglia detected by immunohistochemistry (Stables et al., 2022). The profiles of individual microglia-specific transcripts (**Figure 1B**) suggest a subtle change in microglial phenotype with age, notably down-regulation of *P2ry12* relative to other markers but unlike the reported microglial dyshomeostasis in *Csf1r*^+/-^ mice (Arreola et al., 2021) there was no relationship to *Csf1r* genotype. Furthermore, *Csf2*, reported to drive microgliosis in *Csf1r^+/-^*mice (Chitu et al., 2020), was barely detectable. The most recent report on the *Csf1r*^+/-^ model described an increase in *Trem2* mRNA and evidence that TREM2 participates in demyelination (Biundo et al., 2023a), but both *Trem2* and *Tyrobp* (encoding the signal transducer DAP12) were reduced equally in hippocampus and striatum of *Csf1r*^E631K/+^ mice at both ages (**Table S1**).

**Figure 1.**
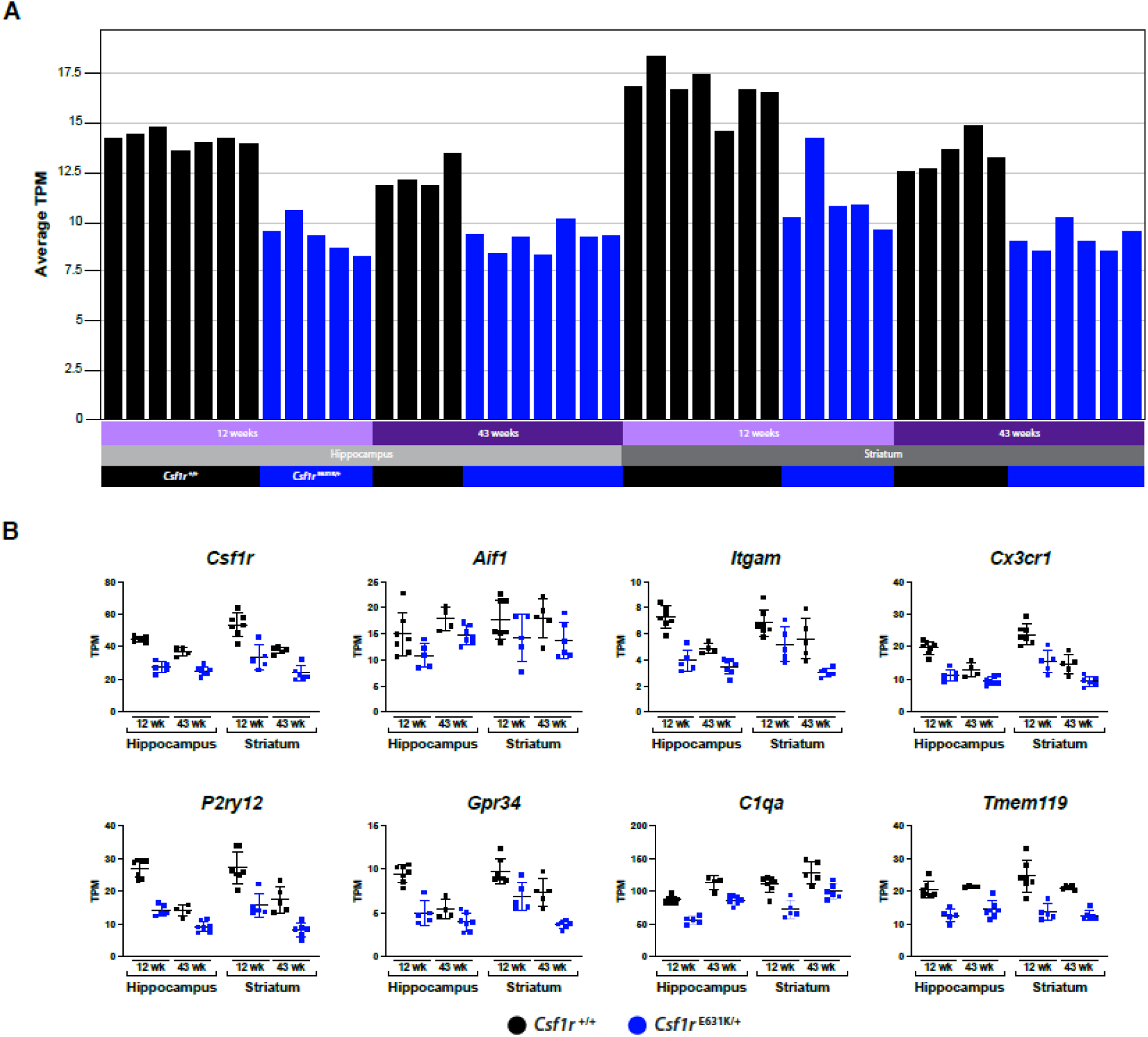
The effect of heterozygous *Csf1r*^E631K^ mutation on gene expression in the hippocampus and striatum. RNA-Seq analysis was performed on hippocampus and striatum from *Csf1r*^+/+^ and *Csf1r*^E631K/+^ mice at 12 and 43 weeks of age (n = 4-7/group). Gene-to-gene co-expression cluster analysis was conducted using Graphia as described in Materials and Methods. Clusters were generated at *r* > 0.8 and MCL inflation value 1.5. The full data set and cluster lists are provided in Table S1A and S1B. (A) Average expression profile of co-expressed genes in cluster 13 (36 genes). X-axis shows the individual samples with columns coloured by group group (black – *Csf1r*^+/+^; teal -*Csf1r*^E631K/+^); Y-axis shows average expression of all genes contained in the cluster in TPM. (B) Gene expression profiles for individual selected genes. Y-axis shows the expression level in TPM. Significance values are not shown on the plots. The full list of pairwise expression comparisons based upon *Csf1r* genotype can be found in Table S4.

The expression analysis did not reveal significant changes in abundance of key genes such as parvalbumin (*Pvalb*) aggrecan (*Acan*), synaptophysin (*Syp*) or bassoon (*Bass*) encoding proteins with reduced expression in *Csf1r*^+/-^ mice (Arreola et al., 2021). Unlike microglia-specific transcripts, markers of brain-associated macrophages, including *Cd163, Mrc1, Lyve1* and *Cd74* were reduced only in the hippocampus at 12 weeks and normalised by 43 weeks (**Table S1A**). **Cluster 1** includes transcripts associated with the vasculature and pericytes (e.g. *Sox17/18, Flt1, Esam, Cdh5, Pecam1, Nos3, Pdgfb Pdgfrb, Ptprb*) each reduced by >50% in both hippocampus and striatum at 43 weeks, consistent with published analysis (Chen et al., 2020) as well as *Dcx, Neurod1, Ncam2* and *Nes* likely reflecting reduced neurogenesis with age (Yousef et al., 2019). However, in neither brain region was there any evidence of an effect of *Csf1r*^E631K/+^ genotype on region-specific or age-dependent changes in gene expression. Notably, there was no significant increase in *Serpina3n* or *C4b*, markers of oligodendrocyte dysregulation in microglia-deficient mice (McNamara et al, 2023). We conclude that aside from reduced microglial density, the dominant *Csf1r* mutation has no significant effect on microglial gene expression, brain development or function. This conclusion is consistent with the previous failure to detect any effect of the heterozygous mutation in behavioral or motor-related assays, brain architecture or pathology (Stables et al., 2022).

### Epistatic interaction between *Csf1r*^E631K/+^ and *Cx3cr1*^EGFP/+^ mutations

The *Cx3cr1-*EGFP line generated by Jung *et al*. (Jung et al., 2000) has been widely used in studies and macrophage and microglial biology. A common assumption in the use of these mice, and subsequent use of a *Cx3cr1-*cre knock-in (e.g. (Arreola et al., 2021)) is that heterozygous *Cx3cr1* loss of function arising from the EGFP insertion has no phenotypic impact. However, Rogers *et al*. (Rogers et al., 2011), in a study of homozygous mutation of the receptor on hippocampal development and plasticity, reported an intermediate phenotype in heterozygotes and Gyoneva *et al*. (Gyoneva et al., 2019) described a transcriptional signature of premature aging in *Cx3cr1*^EGFP/+^ microglia*. Cx3cr1* is expressed exclusively in microglia and homozygous mutation impacts multiple aspects of brain development, including synaptic plasticity and glutamatergic neurotransmission (Basilico et al., 2022; Basilico et al., 2019; Lauro et al., 2008). To test potential epistatic interactions between *Cx3cr1* and *Csf1r* loci in control of brain development, we examined the effect of the *Cx3cr1*-EGFP insertion mutation in *Csf1r*^E631K/+^ mice. In simple terms, we predict that microglia in a double heterozygote will be both reduced in number and dysfunctional.

Compared to WT, *Csf1r*^E631K/+^ mice have around 40% fewer microglia (Stables et al. 2022). **Figure 2A** and **B** show flow cytometry analysis of microglia isolated from WT or *Csf1r*^E631K/+^ mice on the heterozygous *Cx3cr1*-EGFP background using conventional markers CD45 and CD11b. The reduction in microglial abundance in *Csf1r*^E631K/+^ mice is compatible with previous observations and confirms that neither mutation alters the relative expression of the two markers. By contrast to reduced CSF1R protein expression observed in peripheral macrophage populations (Stables et al. 2022) the *Csf1r*^E631K^ mutation did not alter the levels of cell-surface CD115 (CSF1R) detected on isolated microglia (**Figure 2C**, **D**). To compare microglial number across all four genotypes, brain sections from young adult mice were examined using either *Csf1r-*EGFP or *Cx3cr1-*EGFP as indicators. Throughout the brain, microglial cell bodies appeared less dense in *Csf1r*^E631K/+^ compared to wild-type. **Figure 2E** shows representative images of microglia in striatum. Quantitation in cortex (**Figure 2F**) and hippocampus (**Figure 2G**) confirmed a significant reduction in detection of *Cx3cr1*-EGFP^+^ cells in *Csf1r*^E631K/+^ brains.

**Figure 2.**
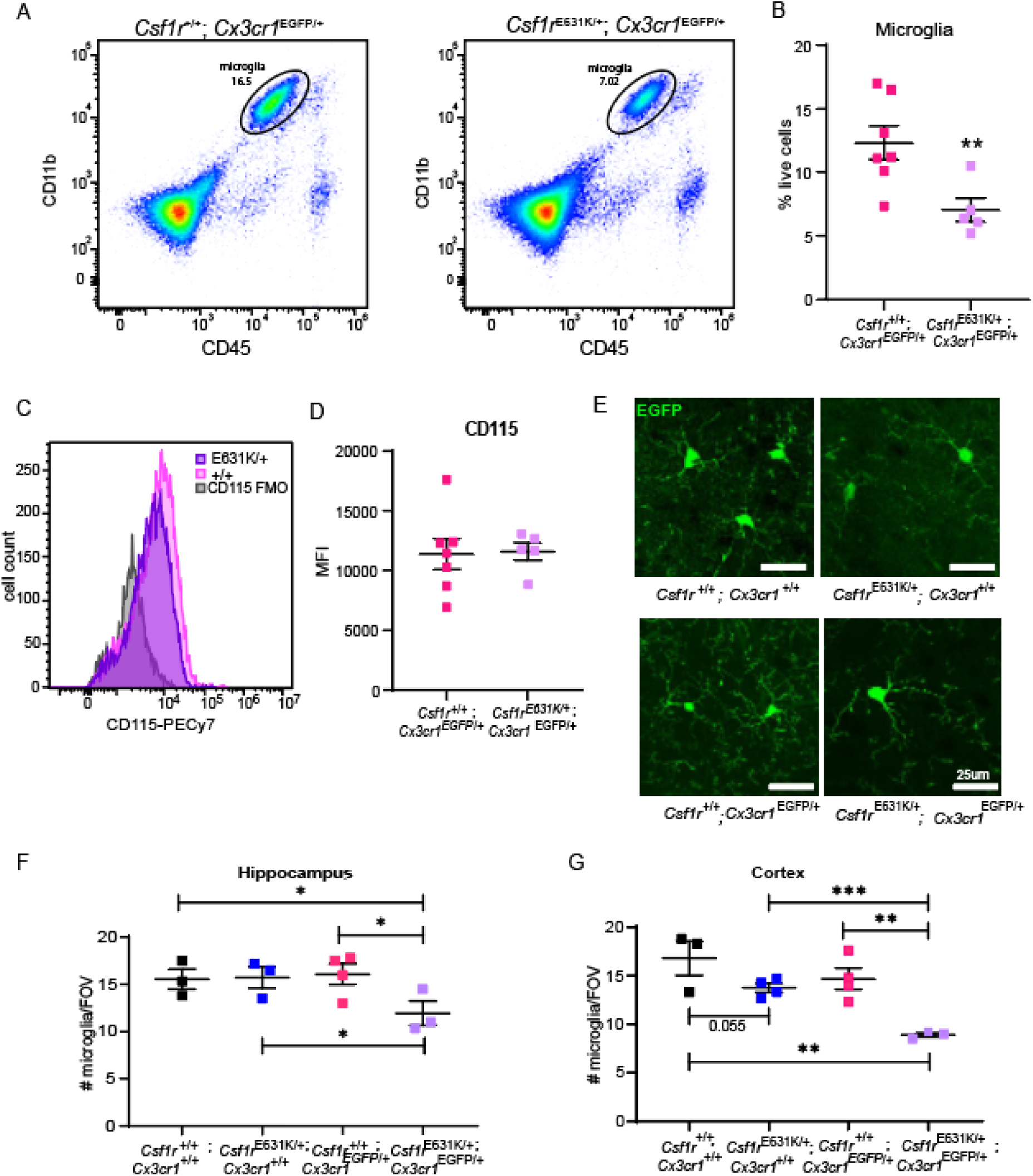
The effect of heterozygous *Cx3cr1* mutation on *Csf1r*^E631K/+^ microglia. (A) Representative flow cytometry plots of microglia in dissociated brain tissue from *Csf1r*^+/+^; *Cx3cr1*^EGFP/+^ and *Csf1r*^E631K/+^; *Cx3cr1*^EGFP/+^ mice. (B) Quantitative analysis of microglia as a % of all live cells. (C) Overlaid, representative histograms for CD115 staining of microglia for each genotype and FMO control. (D) Quantification of CD115 mean fluorescence intensity (MFI). n = 5-7/group. (E) Representative images of GFP^+^ microglia in striatum in *Csf1r*^+/+^; *Cx3cr1*^+/+^, *Csf1r*^E631K/+^; *Cx3cr1*^+/+^, *Csf1r*^+/+^; *Cx3cr1*^EGFP/+^, and *Csf1r*^E631K/+^; *Cx3cr1*^EGFP/+^ mice as indicated. In the upper panels, the reporter is *Csf1r-*EGFP. In the lower panel, the reporter is *Cx3cr1-*EGFP. Scale bar = 25 µm. Microglial density detected with the two reporters, calculated as number of GFP^+^ cell bodies/field of view in the cortex (F) and hippocampus (G). Image analysis is described in Methods. n = 3-4/group. Student’s T Test; * P<0.05, **P<0.01, ***P<0.005.

We next performed RNA-seq analysis on the hippocampus and striatum of adult (12 weeks old) double heterozygotes compared to WT, *Cx3cr1*^EGFP/+^ and *Csf1r*^E631K/+^ controls. The full dataset is shown in **Table S2A;** ranked based upon the expression in double heterozygotes (DHet, heterozygous for both *Csf1r* and *Cx3cr1* mutations) versus WT. As above, the dataset was clustered using the network analysis software, Graphia (Freeman et al., 2022); clusters and their patterns of shared expression (average expression of all transcripts within the cluster) are provided in **Table S2B. Cluster 1,** nearly 4000 transcripts, groups transcripts based upon a shared pattern of down-regulation in both hippocampus and striatum specifically in double heterozygotes (**Figure 3A**), consistent with existence of genetic epistasis. Known microglia-specific transcripts with clustered with *Csf1r* in **Cluster 12.** Analysis of individual microglia-specific transcripts (**Figure 3B**) indicated a complex interaction between the two mutations. Consistent with the reported lack of dosage compensation (Faust et al., 2023), *Cx3cr1* mRNA was reduced by 50% in heterozygous mutant brains and was further reduced by a similar proportion in the double heterozygotes relative to *Csf1r*^E631K/+^. Whereas the heterozygous *Cx3cr1* mutation had no significant effect on the levels of the majority of microglia-specific transcripts in hippocampus or striatum on either WT or *Csf1r*^E631K/+^ genetic background (highlighted in **Table S2A**), a small subset of homeostatic microglia-specific transcripts (*Sall1, Ikzf1, P2ry12, Gpr34, Tgfbr1, Itgam*) was further reduced in parallel with *Cx3cr1* in double heterozygotes (**Figure 3B**, **Table S2A**).

**Figure 3.**
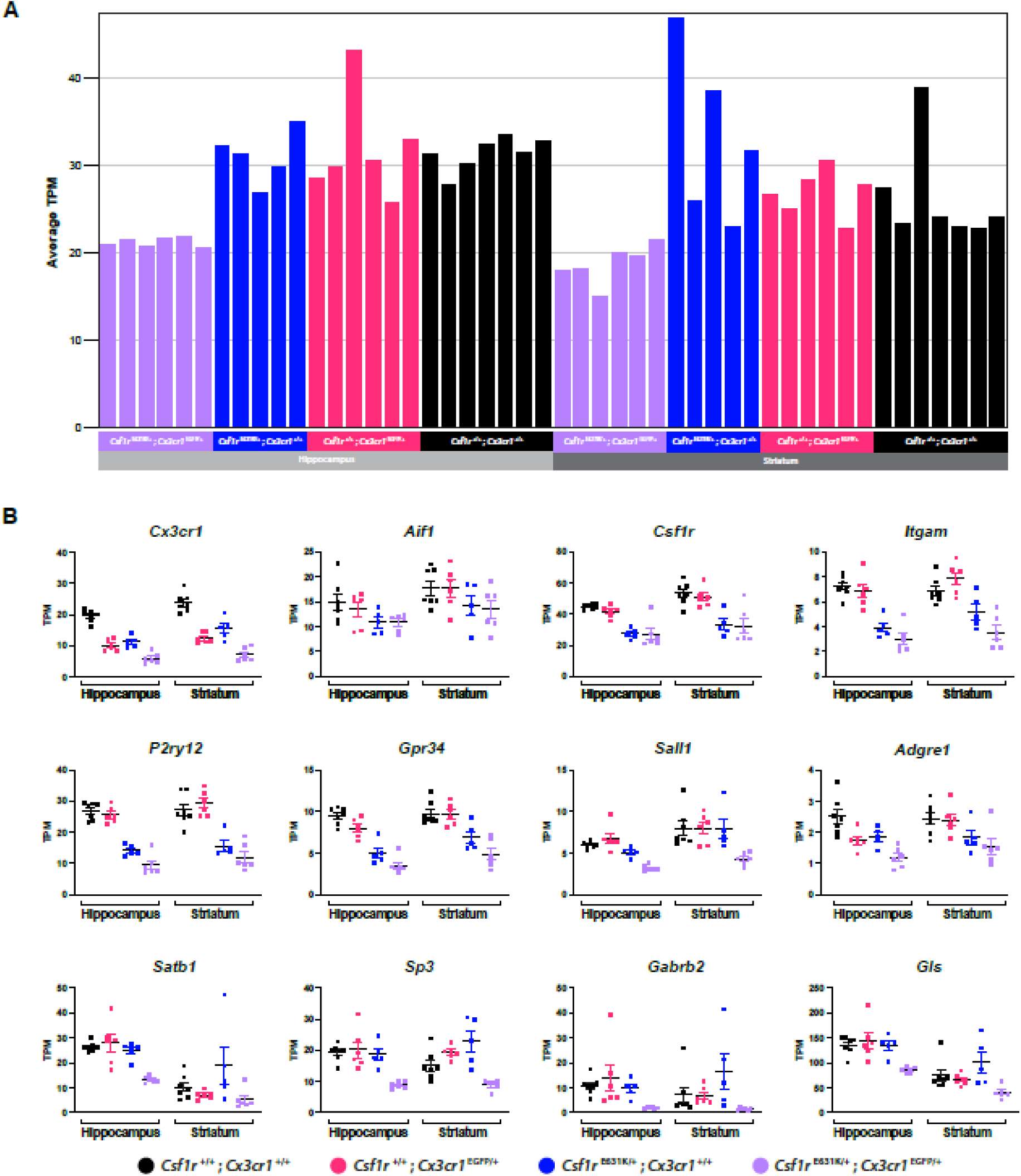
The effect of double heterozygous *Csf1r*^E631K^ and *Cx3cr1*^EGFP^ mutations on gene expression in hippocampus and striatum. RNA-Seq analysis was performed on hippocampus and striatum from *Csf1r*^+/+^; *Cx3cr1*^+/+^, *Csf1r*^E631K/+^; *Cx3cr1*^+/+^, *Csf1r*^+/+^; *Cx3cr1*^EGFP/+^, and *Csf1r*^E631K/+^; *Cx3cr1*^EGFP/+^ mice at 12-15 weeks of age (n = 5-7/group). Gene-to-gene co-expression cluster analysis was conducted using Graphia as described in Materials and Methods. Clusters were generated at *r* > 0.8 and MCL inflation value 2. The full data set and cluster list are provided in **Table S2A** and **S2B**. (A) Average expression profile of co-expressed genes in cluster 1 (3868 genes). X-axis shows the individual samples with columns coloured by group; Y-axis shows average expression of all genes contained in the cluster in TPM. (B) Gene expression profiles for individual selected genes. Y-axis shows the expression level in TPM. Significance values are not shown on the plots. The full list of pairwise comparisons for individual transcripts between controls and double heterozygotes can be found in **Table S4**.

The transcripts in **Cluster 1** share a pattern but the extent of selective reduction of individual transcripts in double heterozygotes varies. Amongst the more abundant transcripts decreased in both regions, we identified subunits of neurotransmitter receptors (e.g *Gabra1/4, Gabrg1, Gabrb2, Gria2, Grik2, Grm1/5*), glutamate transporters (*Slc6a1*), glutaminase (*Gls*) and multiple transcription factors involved in neuronal development (e.g. *Satb1, Sp3, Tcf1l2, Pou3f2, etc*). These are highlighted in **Table S2A** and examples are shown in **Figure 3B**. Conversely, in both hippocampus and striatum, there was a small set of abundant transcripts increased in the double heterozygotes (**Table S2A**) including the metal-binding proteins, *Mt1, Mt2* and *Ptms* as well as *Ptgds*, a regulator of myelination (Pan et al., 2023). Arreola *et al*. (Arreola et al., 2021) identified a set of transcripts up-regulated in the cortex in their *Csf1r*^+/-^ mice, albeit at 8 months of age. We found no significant change in expression of any of these transcripts in either brain region.

Microglia have been attributed many functions in neurogenesis in the hippocampus and CX3CR1 signalling has been implicated (Rogers et al., 2011). A recent study indicated a role for P2RY12 in microglia communication with neurogenic progenitors in the hippocampus (Cserep et al., 2022). In view of the apparent reduction in both *Cx3cr1* and *P2ry12* in double heterozygotes, we examined expression of neurogenic progenitor markers previously identified by single cell RNA-seq (Artegiani et al., 2017). We detected small, but significant reductions in progenitor-enriched transcripts including *Dcx, Tcf4, Neurod1, Homer1, Fgfr3, Sox2, Ncam2, Epha4, Zbtb18, Lrp8, Elavl2/3/4* specifically in *Cx3cr1*^+/-^/*Csf1r*^E631K/+^ hippocampus (**Table S2A**). These data provide strong evidence for a functional interaction between *Csf1r* and *Cx3cr1* and a model for the possible impacts of genetic epistasis between *CSF1R* and other microglia-expressed genes in human neurodegenerative disease.

### The response of Csf1r^E631K/+^ mice to prion infection

Microgliosis is a prominent histopathological characteristic that accompanies the accumulation of disease-specific misfolded host prion protein (PrP^Sc^) in the brain during the development of prion disease (Aguzzi and Zhu, 2017). Microgliosis in prion diseases is associated with increased expression of both *Csf1* and *Il34* (Obst et al., 2017). Based upon the impacts of depletion with CSF1R kinase inhibitors, microglia have been attributed roles in the initiation of prion disease pathology (Carroll et al., 2018; Gomez-Nicola et al., 2013; Race et al., 2022). However, we found that CNS prion disease was accelerated in the complete absence of microglia in *Csf1r*^ΔFIRE/ΔFIRE^ mice without any increase in PrP^sc^ accumulation (Bradford et al., 2022). Infectious prions consist of misfolded isoforms of the host cellular prion protein (PrP^C^). In the present study, the level of *Prnp* mRNA in hippocampus or striatum of *Csf1r*^E631K/+^ mice was unaffected at either age (**Table S1A**) and the expression of PrP^C^ protein in the brain was similarly unaffected (**Figure 4A**,**B**).

**Figure 4.**
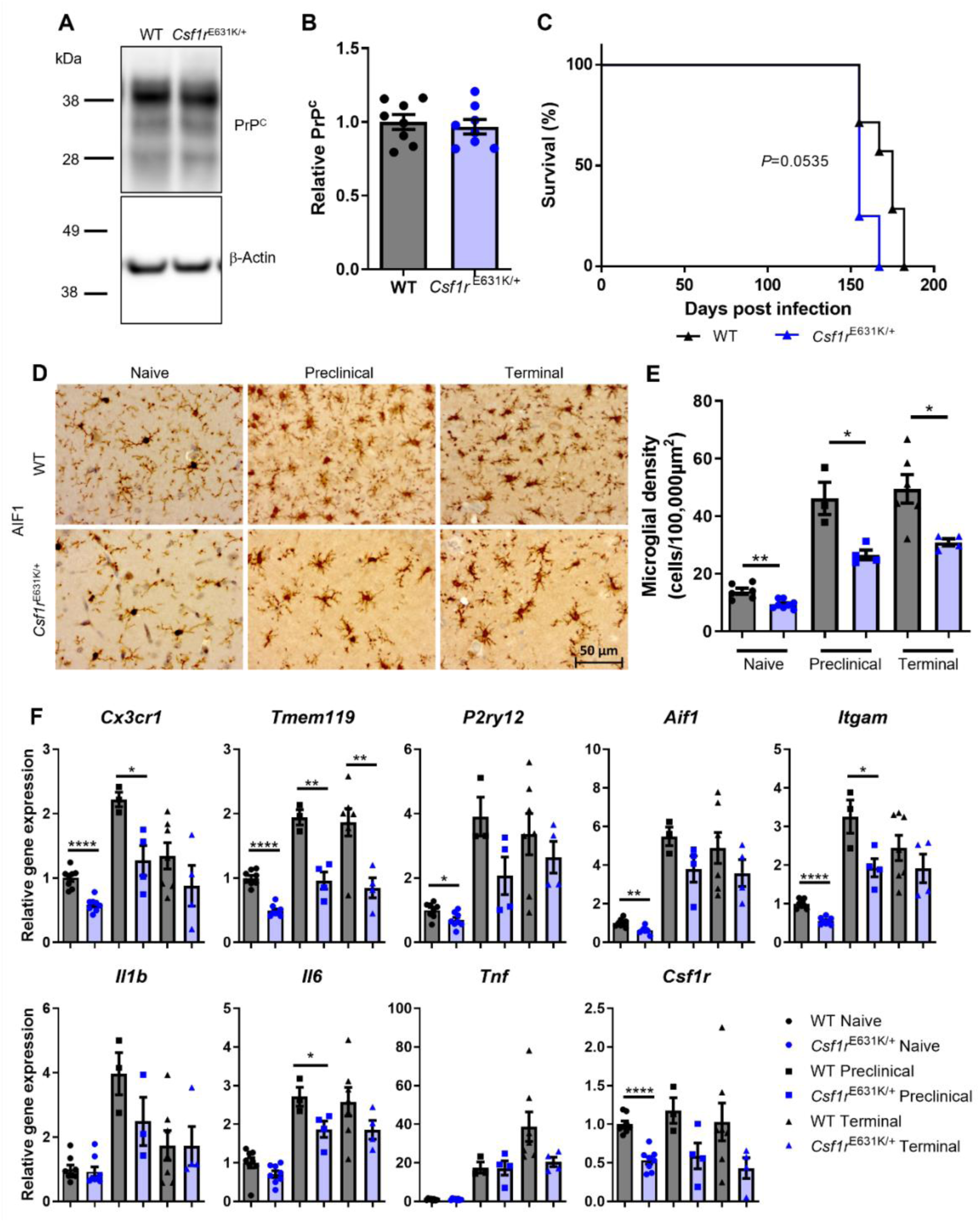
The effect of heterozygous *Csf1r*^E631K^ mutation on the response to prion infection. (A) Western blot detection of PrP^C^ and loading control β-actin in mouse half brains (representative). (B) Densitometry quantitation of the relative abundance of PrP^C^ protein in brains from WT mice and *Csf1r*^E631K/+^ mice. The relative abundance of PrP^C^ protein in each sample was normalized to β-actin and compared to the mean value in the brains of WT mice. Points represent individual mice. Ns. Not significantly different, student’s t-test. Horizontal bar, mean. N=8 mice/group. (C) Survival curve analysis following intracerebral prion challenge shows WT mice and *Csf1r*^E631K/+^ mice succumbed to clinical prion disease at similar times. N=4–7 mice/group. Log-rank (Mantel-Cox) test. ns, not significant (D) Representative immunostaining of IBA1^+^ microglia in the CA1 region of the hippocampus of WT and *Csf1r*^E631K/+^ mice. Scale bar 50 µm. (E) Microglial density in the hippocampus of WT mice and *Csf1r*^E631K/+^ mice calculated via image analysis N=4–8 mice/group, Student’s T Test; * P<0.05, **P<0.01. (F) Relative expression level of the microglia-associated transcripts *Cx3cr1*, *Tmem119*, *P2ry12*, *Aif1*, *Itgam*, *Il1b*, *Il6*, *Tnf* and *Csf1r* in brain. Gene expression was analysed by qRT-PCR and normalized to uninfected WT mice. Points represent individual mice. Horizontal bar, mean. N=3-8 mice per group. Students T-test ns = not significant, *P<0.05, **P<0.01, ****P<0.0001.

Groups of *Csf1r*^E631K/+^ mice and WT mice were infected with ME7 scrapie prions directly into the CNS by intracerebral injection (Bradford et al., 2022). The prion-infected *Csf1r*^E631K/+^ mice succumbed to clinical prion disease at similar times to WT mice (*Csf1r*^E631K/+^ mice 158 ± 6 days, WT mice 170 ± 12 days, respectively, *P* = 0.0545, n = 4-7 mice/group; **Figure 4C**). Immunostaining for IBA1 confirmed that prion infection was accompanied by a 3-fold increase in the density of microglia in the brains of WT mice at 140 dpi and at the terminal stage of disease. In the *Csf1r*^E631K/+^ mice, microglial density was reduced in baseline, preclinical and terminal phases and residual IBA1^+^ cells appeared less ramified (**Figure 4D**, **4E**). The histochemical evidence of altered microglial density was supported by the reduced detection of microglia-expressed transcripts (*Cx3cr1, Tmem119, P2ry12; Aif1, Itgam;* **Figure 4F**) and encoding the disease-induced pro-inflammatory cytokines (*I1b, Il6, Tnf*) (**Figure 4F**). Interestingly, *Csf1r* mRNA was not substantially increased in disease in either WT or *Csf1r*^E631K/+^ mice; likely reflecting the regulation of the gene in response to ligand stimulation (Yue et al., 1993).

Others have suggested that microglia help to protect the brain against prion disease by phagocytosing and sequestering prions (Carroll et al., 2018) but previous analysis of microglia-deficient mice did not support this view (Bradford et al., 2022). Western blot analyses (**Figure 5A**, **5B**) showed that the accumulation of PrP^Sc^ in the brain was unaffected in infected *Csf1r*^E631K/+^ mice compared to WT despite the relative reduction in microglial density. The magnitude and distribution of the neuropathology (vacuolation; **Figure 5C**,**D**) and reactive astrocytosis (**Figure 5C**,**E**) was also indistinguishable in the brains of the clinically-affected *Csf1r*^E631K/+^ mice and WT mice throughout the disease course. One of the impacts of prion disease is extensive demyelination, also a feature of ALSP and the *Csf1r*^+/-^ mouse model (Biundo et al., 2023a). **Figures 5F** and **5G** show the extensive demyelination associated with terminal prion disease which was neither prevented, nor exacerbated, in the *Csf1r*^E631K/+^ mice. Notably, the uninfected WT and *Csf1r*^E631K/+^ mice were also indistinguishable at this age (6-8 months).

**Figure 5.**
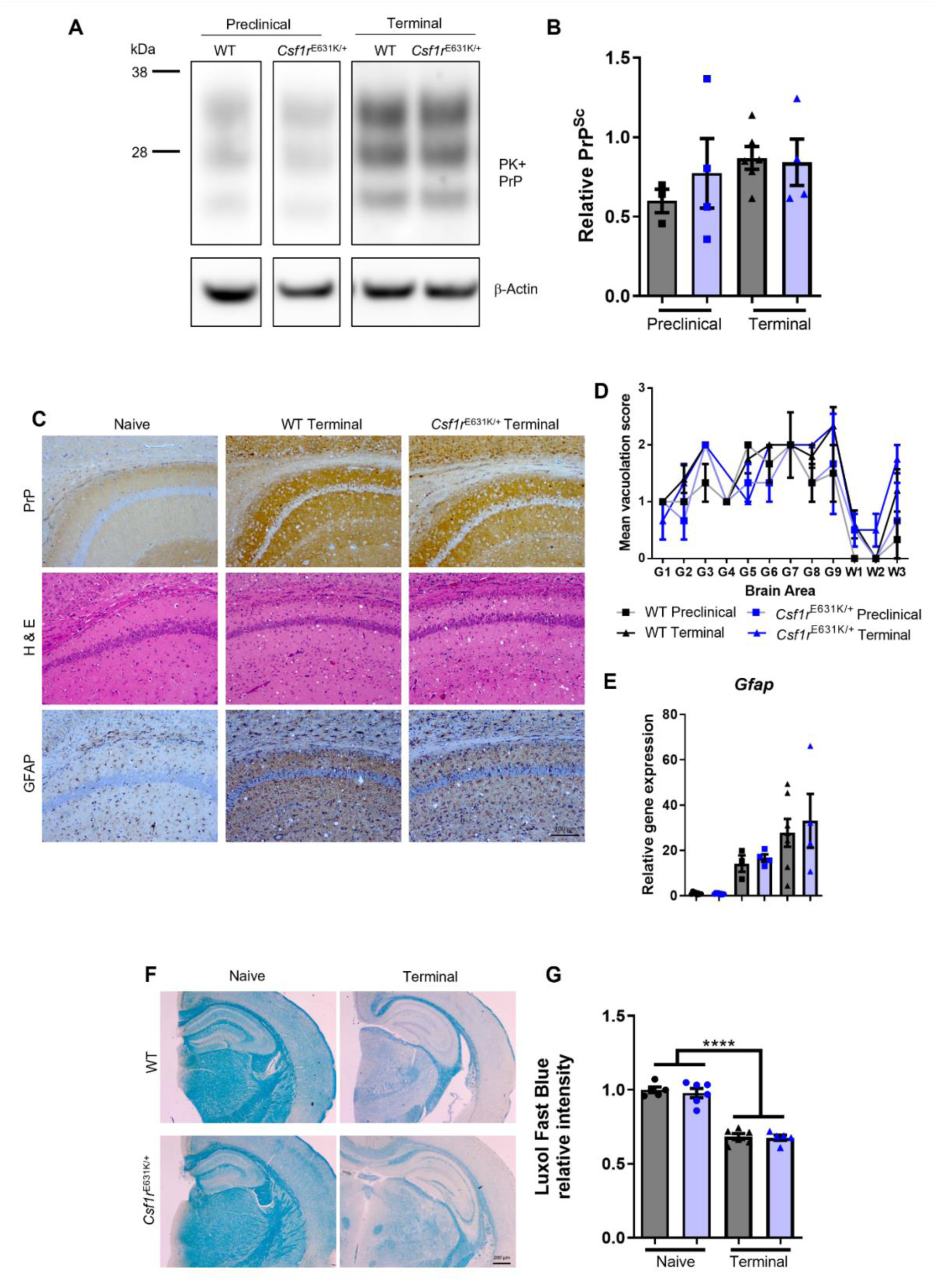
The effect of heterozygous *Csf1r*^E631K^ mutation on neuropathology associated with prion infection. (A) Western blot detection of proteinase-K resistant prions in preclinical and terminal prion infected brains. (B) Quantification of the relevant amounts of PrP^Sc^ prions by densitometry, normalised by β-actin loading control and expressed relative to native PrP^C^ abundance in naïve brain. Points represent individual mice, horizontal bar = mean. Students T-test, ns = not significant. (C) Representative immuno-or histo-chemical staining reveals similar prion accumulation (PrP^Sc^), prion-specific vacuolation (Hematoxylin and Eosin) and astrocyte reactivity (GFAP) in terminal vs naïve brains of WT mice and *Csf1r*^E631K/+^ mice. (D) Severity and distribution of the vacuolar pathology in the brains of prion-infected WT mice and *Csf1r*^E631K/+^ mice. Vacuolation scored on H&E sections of each brain in the following 9 grey (G) matter and 3 white (W) matter brain regions: G1, dorsal medulla; G2, cerebellar cortex; G3, superior colliculus; G4, hypothalamus; G5, thalamus; G6, hippocampus; G7, septum; G8, retrosplenial and adjacent motor cortex; G9, cingulate and adjacent motor cortex; W1, inferior and middle cerebellar peduncles; W2, decussation of superior cerebellar peduncles; and W3, cerebellar peduncles. Groups as indicated. (E) Relative expression level of *Gfap* via qRT-PCR. N=3-8 mice per group. Students T-test ns = not significant. (F) Representative images of luxol fast blue staining of age-matched naïve and terminal prion infected WT mice and *Csf1r*^E631K/+^ mice. (G) Quantitation of luxol fast blue intensity. Points represent individual mice. Horizontal bars, mean. N=5-6 mice per group. ANOVA with Tukey’s multiple comparisons. **** P<0.0001.

### The response of Csf1r^E631K/+^ mice to experimental autoimmune encephalitis (EAE)

The myeloid cell population of the brain is also increased in EAE, a widely-studied model of multiple sclerosis, due to both proliferative expansion of the microglial cell population and recruitment of blood monocytes (Ajami et al., 2011; Hagan et al., 2020; Hwang et al., 2022). Several recent reports using CSF1R kinase inhibitors have implicated inflammatory microglia and monocytes in demyelination in EAE (Hwang et al., 2022; Montilla et al., 2023; Nissen et al., 2018). We therefore tested the impact of the *Csf1r*^E631K/+^ genotype in this experimental model (**Figure 6A**). EAE was initiated by immunisation with myelin oligodendrocyte glycoprotein (MOG35-55) and clinical scores were assessed as described (Bittner et al., 2014; Khan et al., 2014). In view of the known sex differences in susceptibility (Bourel et al., 2021; Wiedrick et al., 2021) this model was studied in females. For this experiment, the *Csf1r*-EGFP reporter (Sasmono et al., 2003) enabled visualisation of infiltrating myeloid cells.

**Figure 6.**
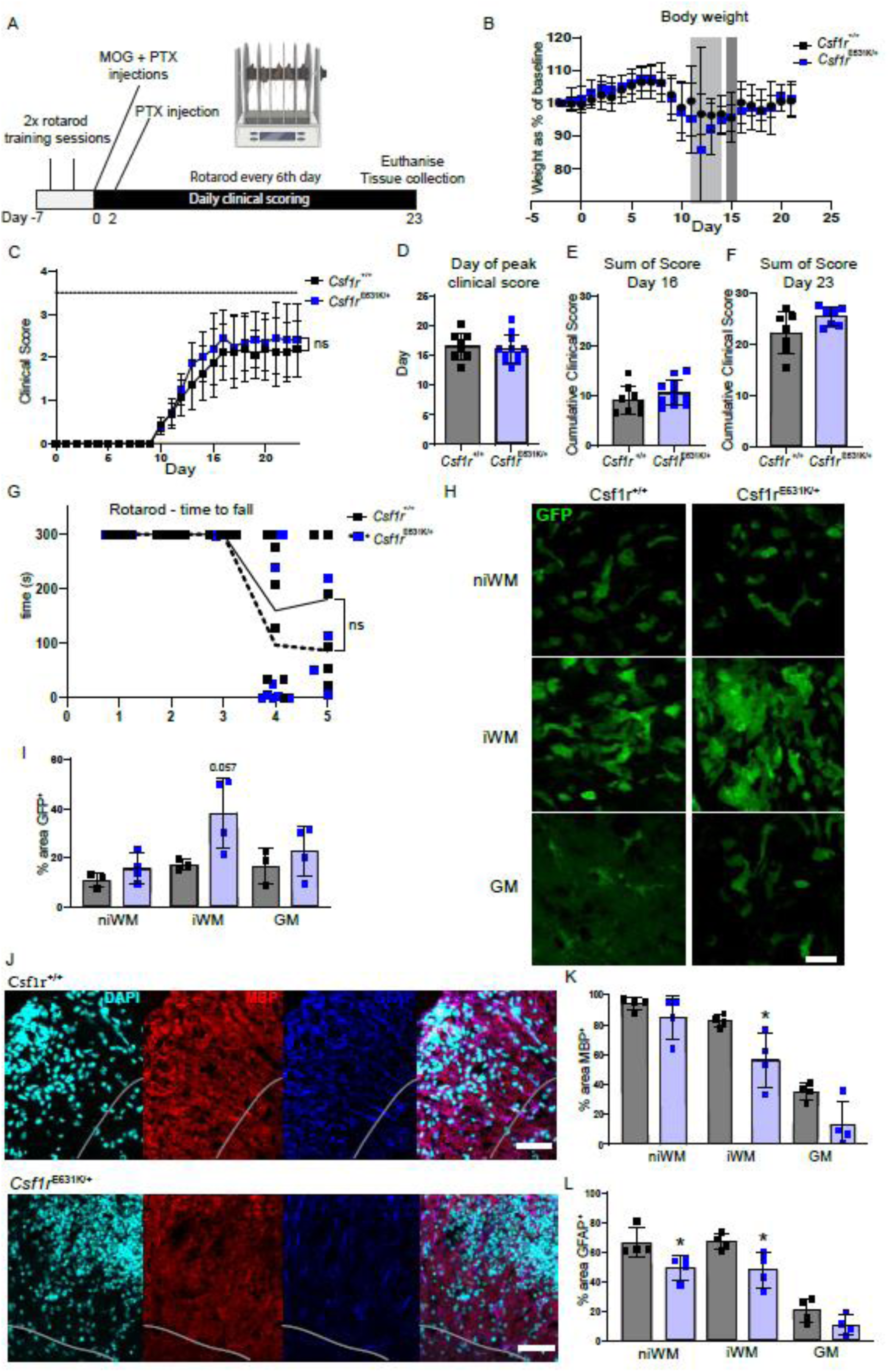
The effect of heterozygous *Csf1r*^E631K^ mutation on disease progression in EAE. (A) The timeline for EAE induction, clinical scoring, and rotarod assessment. Mice were euthanised on day 23 post-EAE induction, or when they reached a clinical score of 3.5 (hindlimb paralysis present with signs of forelimb weakness). (B) The baseline weight of each mouse was taken 2 days prior to EAE induction and body weight changes were calculated daily. n=8-11 female mice/group. Light grey box indicates that weight as a % of the baseline body weight was significantly reduced in heterozygous mice, and dark grey in WT mice only. (C) The mean clinical scores for mice of both genotypes from day 0 – 23. The day at which clinical score peaked (D), sum of daily clinical scores at this peak (day 16 -E), and for mice that survived to the endpoint, day 23 (F). (G) The time that each mouse remained on the accelerating rotarod was measured up to a maximum of 300s. n=8-11/group. (H) *Csf1r*-EGFP imaging in the spinal cord. Scale bar = 20 µm. The non-infiltrated white matter (niWM), infiltrated white matter (iWM), and the grey matter were assessed in 3 spinal cord sections (3 regions per section) per animal for the GFP+ % area (I). n = 3-4/group. (J) Representative MMP and GFAP immunofluorescence staining in the spinal cord. Grey line delineates WM from GM. Scale bar = 50µm. The MBP+ (K) and GFAP+ (L) staining area (%) was calculated in 3-4 sections per animal. n = 4/group. * p<0.05 Mann-Whitney test.

Aside from a marginal acceleration of body weight loss, there were no significant differences in disease pathology between WT and *Csf1r*^E631K/+^ mice (**Figure 6 B-G**). In EAE, *Csf1r*-EGFP^+^ myeloid cells were selectively increased in the involved white matter in the spinal cord irrespective of genotype (**Figure 6 H**,**I**). However, immunolocalisation of myelin basic protein (MBP) indicated a greater reduction in the MBP in the involved white matter of *Csf1r*^E631K/+^ compared to control mice (**Figure 6** **J,K**). Microglia-astrocyte interaction has been strongly implicated in the pathology of multiple sclerosis and EAE (Absinta et al., 2021). Based upon quantitative analysis of GFAP, disease-associated astrocytosis in the spinal cord white matter was significantly reduced in the *Csf1r*^E631K/+^ mice compared to WT (**Figure 6L**).

Although the spinal cord is the key location of disease-associated demyelination underlying motor dysfunction, microgliosis and microglial activation contributes to neuropathology throughout the brain (Bourel et al., 2021; Diebold et al., 2023). To assess the effect of the *Csf1r*^E631K/+^ mutation on this response, we performed total RNA-seq profiling of hippocampus and striatum of mice of each genotype with a similar clinical score of 2 (hind limb weakness/partial paralysis of hindlimbs). The results are shown in **Table S3.** The gene lists in **Table S3A** are ranked based upon the ratio of expression in disease (EAE) versus control in the WT cohort. **Table S3B** shows the co-regulated clusters identified by network analysis. By contrast to previous reports (Hagan et al., 2020; Hwang et al., 2022) neither *Csf1*, nor *Il34* mRNA was increased in EAE in either brain region and both *Csf2* and *Csf3* remained undetectable. A co-regulated cluster of 400 transcripts (**Cluster 4**) includes the majority of microglia-specific transcripts identified above. The average profile is shown in **Figure 7A** and expression of selected individual transcripts in **Figure 7B**. In broad terms, the increased detection of microglial markers (*Csf1r, Aif1, Adgre1, Itgam*) is consistent with >2-fold increase in microglia in the hippocampus and somewhat less in striatum in response to EAE. The fold-increase for microglia-related transcripts was not significantly different between *Csf1r*^E631K/+^ mice and WT, so the absolute abundance of all of the EAE responsive transcripts in **Cluster 4** was reduced in both brain regions in the mutants. However, in both brain regions, disease was associated with a much greater increase in detection of transcripts encoding class II MHC-related (e.g *Cd74, H2-Aa*), effectors of the interferon response, inflammatory chemokines and complement components (*C3, C1q, C4b*) which also formed part of **Cluster 4**. Complement component C3, identified as a key mediator of hippocampal neurodegeneration in EAE (Bourel et al., 2021) was increased >20-fold in hippocampus. Increased expression of C1Q has been implicated in astrocyte activation in EAE (Absinta et al., 2021). The astrocyte marker *Gfap* was increased at least 2-fold in both brain regions (**Figure 7B**) and formed part of **Cluster 14,** containing genes which share relatively elevated expression in hippocampus. The expression of other transcripts in **Cluster 14** was variable between replicates so the effect of the *Csf1r* mutation was not clear. On the other hand, genes associated with homeostatic microglial function, including *Sall1, Cx3cr1, Gpr34, P2ry12/13*, *Slc2a5, Tmem119* were either unaffected or reduced (see *P2ry12* in **Figure 7B**) in response to EAE, again independent of *Csf1r* genotype. EAE has also been associated with functional alterations in brain endothelial cells and the blood brain barrier (Munji et al., 2019). A recent single cell RNA-seq analysis (Fournier et al., 2023) of EAE brain attributed some of the inducible genes in **Cluster 4**, notably *Lcn1*, to endothelial cell activation. The presence of *Lcn1* within **Cluster 4** suggests that it is actually primarily expressed by microglia. Munji *et al* (Munji et al., 2019) identified around 130 transcripts that were up-regulated in endothelial cells isolated from EAE brain at different disease stages. We did not detect a significant increase in any of these markers. However, **Cluster 17** (**Table 3B**) groups transcripts that were down-regulated in EAE in both hippocampus and striatum regardless of *Csf1r* genotype. This Cluster contains *Pecam1, Cdh5, Sox17* (**Figure 7B**) and many other endothelial cell markers, suggesting either endothelial loss or dedifferentiation.

**Figure 7.**
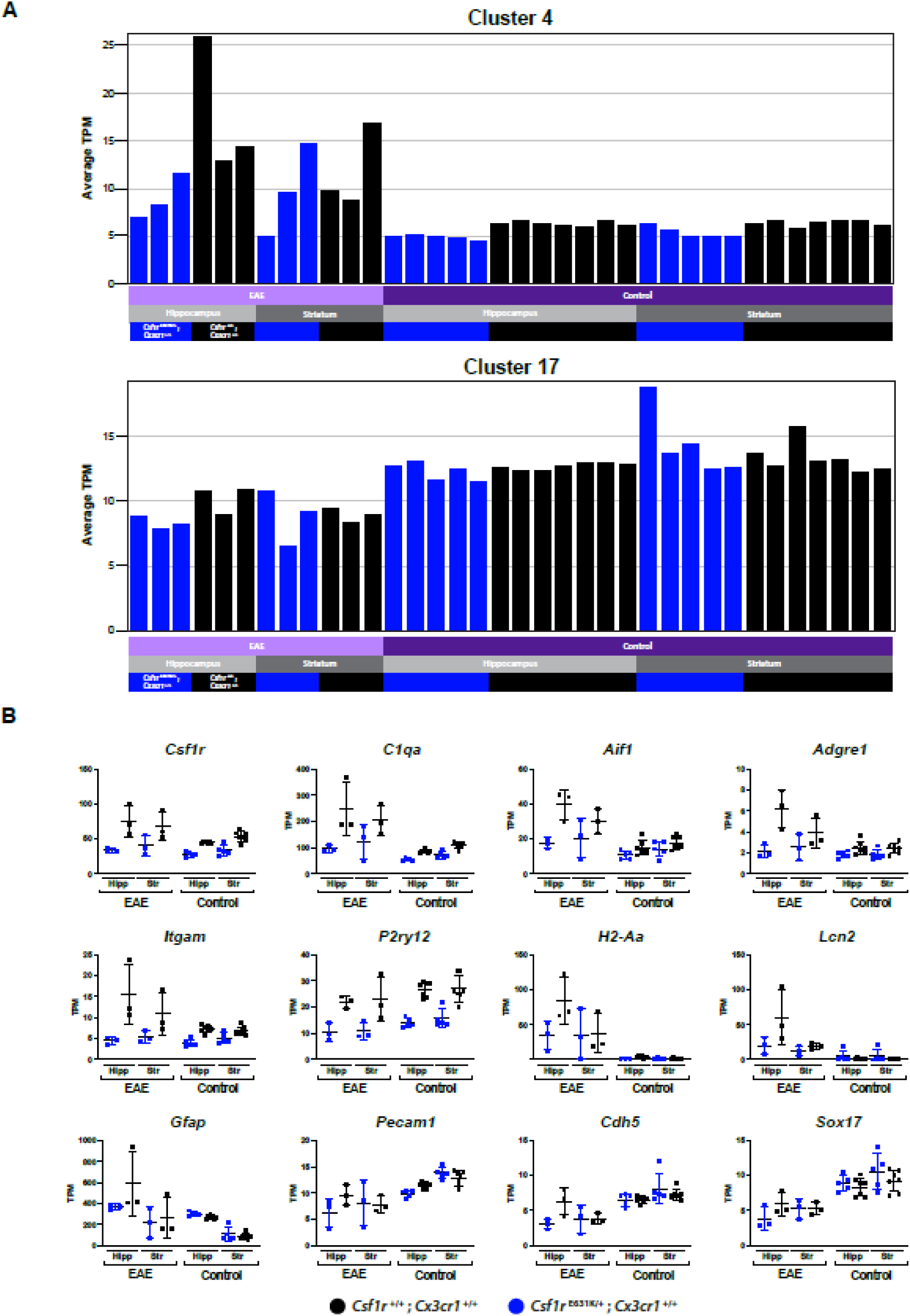
Comparative analysis of changes in gene expression associated with EAE in *Csf1r*^+/+^ and *Csf1r*^E631K/+^ mice. RNA-Seq analysis was performed on hippocampus and striatum from *Csf1r*^+/+^ and *Csf1r*^E631K/+^ mice with or without EAE at 12-15 weeks of age (n = 3-7/group). All of the mice in the EAE group had a clinical score of 2 (hind limb weakness/partial paralysis of hindlimbs). Gene-to-gene co-expression cluster analysis was conducted using Graphia as described in Materials and Methods. Clusters were generated at *r* > 0.8 and MCL inflation value 2. The full data set and cluster list are provided in Table S3A and S3B. (A) Average expression profile of co-expressed genes in Clusters 4 (400 genes) and 17 (49 genes). X-axis shows the individual samples with columns coloured by group; Y-axis shows average expression of all genes contained in the cluster in TPM. (B) Gene expression profiles for individual selected genes. Y-axis shows the expression level in TPM. Significance values are not shown on the plots. The full list of pairwise comparisons can be found in Table S4.

## Discussion

This study has addressed two related issues, the molecular basis for leukoencephalopathy associated with *CSF1R* mutations and the quantitative importance of CSF1R signalling in brain homeostasis. Several recent studies have described a mild microgliosis and consequent neuropathology in mice with a heterozygous null *Csf1r* mutation (Arreola et al., 2021; Biundo et al., 2020; Biundo et al., 2023b; Chitu et al., 2020; Chitu et al., 2015). On this basis, a haploinsufficiency mechanism for the human disease has been proposed (Chitu et al., 2021). It is not yet clear that haploinsufficiency is sufficient to cause the human disease (Chitu et al., 2021; Hume et al., 2020). As an alternative model to inbred mice, we detected no microglial phenotype or neuropathology in aged *Csf1r*^+/-^ rats (Patkar et al., 2021). The loss of microglia, rather than microgliosis, appears to be the hallmark of the human disease (Berdowski et al., 2022; McNamara et al., 2023; Oosterhof et al., 2018).

Regardless of whether a dominant-negative model is accepted, the *Csf1r*^E631K/+^ mouse is certainly heterozygous for a mutation that abolishes CSF1-dependent cell survival when expressed in a growth factor-dependent cell line (Pridans et al., 2013) and phenocopies a complete knockout when bred to homozygosity(Stables et al., 2022). The phenotype is quite clearly distinct from *Csf1r^+/-^* models studied by others (Chitu et al., 2021). In common with te effect of similar heterozygous *Csf1r* coding mutations in zebrafish and in patient brain (Berdowski et al., 2022) we see a consistent reduction in microglial density, supported by the loss of microglia-specific transcripts in RNA-seq analysis, rather than microgliosis. There are several possible explanations for the different phenotypes reported for heterozygous *Csf1r* mutation in mice; environment, genetic background and genotype, including the nature of the mutation. The original *Csf1r* null mutation retains the targeting vector and was not generated on a C57BL/6J background (Dai et al., 2002). In theory, the targeted allele could influence expression of neighboring genes (notably *Pdgfrb,* which contains upstream regulatory elements controlling *Csf1r* (Rojo et al., 2019)) and despite extensive backcross, could also retain linked alleles from the parent 129 strain. One argument against that suggestion is that microgliosis and neuropathology was also observed in a conditional heterozygous mutation (*Csf1r*^fl/+;^ *Cx3cr1*^Cre/+^) (Arreola et al., 2021; Biundo et al., 2020). However, that model is actually a double heterozygote for *Csf1r* and *Cx3cr1* mutations (see below) and complicated further by the known genotoxic effects of expression of Cre recombinase (Faust et al., 2023; Hume, 2023). Chitu *et al*. (Chitu et al., 2021) reported that progression of neuropathology in the *Csf1r*^+/-^ model was accelerated by a high fat breeder diet and there is substantial literature linking dietary fat and microglial function (e.g. (Kim et al., 2019)). Interestingly, *Csf1r*^ΔFIRE/+^ mice on normal chow also showed evidence of microgliosis when analysed at UC Irvine (Stables et al., 2022), the same facility as Arreola *et al*. (Arreola et al., 2021) but the same line in Edinburgh had reduced microglia in the embryo {Munro, 2020 #254).

In the heterozygous null models, microgliosis has been associated with increased expression of alternative microglial growth factors, *Csf2* and *Csf3* {Chitu, 2020 #194;Biundo, 2023 #337}. These transcripts were not induced in *Csf1r*^E631K/+^ brain, even in the EAE model (**Tables S1A, S3A**) and were also not detected in ALSP patient brains (Berdowski et al., 2022). *Csf2* is highly-expressed in innate lymphoid cells (ILC) associated with mucosal surfaces (Forrest et al., 2014; Jacquelot et al., 2022). Since CSF2 crosses the blood brain barrier (McLay et al., 1997) it is conceivable that *Csf1r*^+/-^ mice have an altered response to peripheral stimuli including infectious agents. Whatever the explanation for microgliosis in *Csf1r*^+/-^ mice, our data and that of others (Berdowski et al., 2022) provides additional support for a dominant-negative model for the impact of kinase-dead CSF1R mutations leading to reduced microglial numbers and loss of their trophic functions. This model is compatible with more severe impacts of complete loss of microglia and homozygous *Csf1r* mutations (Chadarevian et al., 2024; Chitu et al., 2021; Hume et al., 2020; McNamara et al., 2023; Munro et al., 2024).The distinction is clearly crucial, since we would argue that treatment with a CSF1R kinase inhibitor (Arreola et al., 2021) would be likely to accelerate rather than inhibit disease progression in patients.

The *Csf1r*^+/-^ model of neuropathology depends upon the C57Bl/6J inbred mouse genetic background. Compared to other strains, C57BL/6J mice have a wide range of genetic variants that impact macrophage function including a coding variant in the *Csf1r* gene (Hume, 2023). The *Csf1r*^+/-^ brain phenotype includes ventricular enlargement (Chitu et al., 2015). C57BL/6J mice exhibit strain-specific susceptibility to development of hydrocephalus in response to a diverse array of mutations affecting ependymal cells (Hume, 2023). Microglial activation could potentially be a downstream consequence of hydrocephalus (Brown et al., 2023). We see no evidence of ventricular enlargement in the *Csf1r*^E631K/+^ brains even at 15 months of age (Stables et al., 2022). Whether the difference between the *Csf1r*^E631K/+^ and *Csf1r^+/-^* models is due to a subtle difference in mouse genetic background cannot be determined. Here we have addressed the possible importance of genetic background by examining the interaction between heterozygous *Csf1r* and *Cx3cr1* mutations. This is analogous in some respects to the conditional heterozygous knockout of *Csf1r* described by others (Arreola et al., 2021; Biundo et al., 2020). By contrast to the report by Gyoneva *et al*. (Gyoneva et al., 2019) the heterozygous *Cx3cr1* mutation alone had no effect apart from the expected 50% reduction in *Cx3cr1* mRNA (Faust et al., 2023). In the double heterozygotes a small subset of “homeostatic” microglia-specific transcripts (*Sall1, Ikzf1, P2ry12, Gpr34, Tgfbr1, Itgam*) was further down-regulated. Hence, microglial dyshomeostasis reported in conditional heterozygous *Csf1r* mutation likely depends upon the *Cx3cr1*-cre driver (Arreola et al., 2021). We also observed a coordinated decrease in expression of transcripts encoding receptors for neurotransmitters and multiple genes involved in glutamate metabolism. These changes in hippocampal and striatal gene expression were not evident in microglia-deficient homozygous *Csf1r*^ΔFIRE^ mice (Rojo et al., 2019) or in mice depleted of microglia using a CSF1R kinase inhibitor (Elmore et al., 2014) where the only significant change was the loss of the microglial signature. CX3CR1/CX3CL1 signaling may regulate postnatal maturation of glutamatergic/GABAergic synapses but the effects of homozygous *Cx3cr1* mutations are relatively small (Basilico et al., 2022; Basilico et al., 2019; Bertot et al., 2019; Corsi et al., 2022) and gene expression has not been reported. The complex interaction between CSF1R and CX3CR1 was evident in a reported decline in spontaneous and evoked glutamatergic activity in chronic PLX5622-treated mice that was prevented in *Cx3cr1^-/-^* mice (Basilico et al., 2022). The effect of the double heterozygote on gene expression is also quite distinct from the *Csf1r*^fl/+;^ *Cx3cr1*^Cre/+^ model (Arreola et al., 2021; Biundo et al., 2020) and the minimal effects of inducible *Cx3cr1^ER2^*^Cre/+^ (Faust et al., 2023). The important conclusion is that there is clear evidence of epistasis. In the context of human disease, the most obvious potential epistatic interaction for *CSF1R* mutations would be with the common variants in the *APOE* gene that regulate microglial function and are associated with susceptibility to dementia (Haney et al., 2024).

Amongst the genes down-regulated specifically in the double heterozygote, *Adnp* (activity-dependent neurotropic protein) was reduced 75-85% in both hippocampus and striatum. ADNP is a transcription factor component of the SWI/SNF chromatin remodeling complex. Mutations in humans are associated with a neurodevelopmental syndrome and haploinsufficiency in mice leads to synaptic plasticity defects, excessive long-term potentiation and impaired learning and memory (Cho et al., 2023). The most down-regulated neurotransmitter receptor gene, *Gabrb2*, encodes the β2 subunit that combines with the α and γ subunits to form the most abundant mammalian subtype of GABA-A receptors, induced during postnatal development. Mutations in this gene are also associated with a wide range of neuropathologies in humans and in mice (Barki and Xue, 2022). Aside from the direct relevance to understanding CSF1R-related encephalopathies, these results demonstrate that the impact of heterozygous loss of CX3CR1 cannot be ignored in models in which *Cx3cr1*-cre is used to drive conditional deletions of genes of interest in microglia (Faust et al., 2023).

We investigated the effect of *Csf1r*^E631K/+^ in two experimental models of neuropathology, prion disease and autoimmune encephalitis. Expression profiling and immunolocalization of microglia was consistent with the role of CSF1R signaling in microgliosis and the dominant-negative model (Stables et al., 2022) in that both basal and increased microglial signatures were repressed. Previous studies in the *Csf1r*^ΔFIRE^ mice described a more rapid onset of pathology in both prion disease (Bradford et al., 2022) and an Alzheimer’s model (Kiani Shabestari et al., 2022). The analysis of the prion model here indicates that CSF1R signaling is required for both basal maintenance of microglia and to generate and sustain the increase in microglial population in response to prion infection (Askew et al., 2017). Since microgliosis is driven by increased CSF1R signals, the result is consistent with the dominant-negative model. However, by contrast to the effect of complete microglial deficiency (Bradford et al., 2022) and treatment with a CSF1R kinase inhibitor (Carroll et al., 2018) the reduction in absolute microglial abundance in the *Csf1r*^E631K/+^ mice was not sufficient to alter the course of disease.

Montilla *et al*. (Montilla et al., 2023) reported a delay in disease onset in EAE when microglia were depleted more completely with PLX5622. Microglial expansion and inflammatory activation in response to EAE was reduced in the *Csf1r*^E631K/+^ mutant mice but this reduction was not sufficient to alter the disease course nor to prevent profound alterations in microglial gene expression within the brain. The relative deficiency of microglia *Csf1r*^E631K/+^ mice may actually exacerbate myelin loss in the spinal cord (**Figure 6**) consistent with their proposed role in preservation of myelin integrity (McNamara et al., 2023).

Taking all of the data together, the primary CSF1R-dependent function of microglia appears to be neuroprotective (Chadarevian et al., 2024; McNamara et al., 2023; Munro et al., 2024). By extension, the loss of that function in individuals with dominant-acting mutations in *CSF1R* likely predisposes to neuropathology triggered by environmental stimuli (e.g. diet, infection, autoimmunity) and/or by interaction with other genetic variants that compromise microglial function including the numerous genetic variants associated with susceptibility to dementia or chronic inflammatory disease.

## Materials and Methods

### Animal breeding

In the original generation of the *Csf1r*^E631K/+^ line (C57BL/6J.Csf1r^Em1Uman^ (Tg16)) donor and recipient mice were C57BL/6JOlaHsd. The progeny were further backcrossed at least 10 times to C57BL/6JCrl. These mice were used for studies of prion disease in Edinburgh. Subsequent to the transfer to Australia, the mice were rederived and bred and maintained in specific pathogen free facilities at the University of Queensland (UQ) facility within Translational Research Institute. To enable visualisation of myeloid populations in tissues, the *Csf1r*^E631K^ line was bred to the *Csf1r*-EGFP reporter transgenic line (Sasmono et al., 2003) also backcrossed >10 times to the C57BL/6JArc genetic background. *Cx3cr1*-EGFP mice on the C57BL6/J background (Jung et al., 2000) were imported and maintained by the University of Queensland Biological Resources. Genotyping of the mouse lines was carried out as described previously (Jung et al., 2000; Sasmono et al., 2003; Stables et al., 2022). All studies in Australia were approved by the Animal Ethics Committee of the University of Queensland. Mice were housed in individually ventilated cages with a 12 h light/dark cycle, and food and water available *ad libitum*. They were bred under specific pathogen free conditions. The *in vivo* mouse studies undertaken in the UK were performed under the authority of a UK Home Office Project Licence and in accordance with the regulations of the UK Home Office ‘Animals (scientific procedures) Act 1986’. Approval to undertake these studies was obtained following review by the University of Edinburgh’s ethical review committee.

### Key reagents

All key reagents including source and catalogue number are provided in Table 1.

### Tissue processing

For immunohistochemistry (IHC) tissues were post-fixed in 4% paraformaldehyde (PFA) for ∼6 h, then transferred to Phosphate Buffered Saline (PBS) with 0.01% sodium azide. A cohort of animals were fixed via transcardiac perfusion using PBS followed by 4% PFA. Tissues were embedded in paraffin and sectioned using core histology facilities at the Roslin Institute, Edinburgh or the Translational Research Institute, Brisbane. Alternatively, tissues were cryoprotected in a 15% then 30% sucrose solution in PBS overnight, then frozen in OCT.

### Immunohistochemistry

For IHC analysis of brain and spinal cord tissue from EAE experimental animals, sections were cut using a Leica RM2245 microtome or a Leica CM1950 cryostat. 12μm frozen sections were placed directly on Superfrost Plus Gold slides and kept at -30⁰C until IHC. 30μm serial sagittal sections were collected from *Cx3cr1*^GFP/+^ animals in a tempero-medial manner (1 in 12 series) using a vibratome (Leica VT 1200S) and kept as free-floating sections in ice-cold PBS. Free-floating brain sections were incubated at room temperature (RT) for 30 min in permeabilization buffer (0.1% Triton-X100 in PBS) followed by 90 min in blocking solution (5% normal goat serum (NGS), 0.3% Triton-X100 in PBS). Sections were then incubated overnight at 4^◦^C under orbital agitation in primary antibody against defined surface markers. Following 3 × 10 min washes in permeabilization buffer, slices were incubated in the appropriate secondary antibody diluted in blocking solution, for 90 min at RT in the dark. Slices were then washed in permeabilization buffer, followed by a 5 min incubation with 4′,6-diamidino-2-phenylindole (DAPI) diluted in PBS, and a further 10 min wash in permeabilization buffer. All sections were washed with PBS for 5 min and mounted with Fluorescence Mounting Medium. Images were acquired on an Olympus FV3000 confocal microscope.

For the prion infection study, paraffin-embedded sections (thickness 6 μm) were deparaffinised and pre-treated by autoclaving in target retrieval solution at 121°C for 15 min. Endogenous peroxidases were quenched by immersion in 4% hydrogen peroxide in methanol or PBS for 10 min. Prior to immunostaining to detect PrP using the prion-specific antibody described previously (McCutcheon et al., 2014), sections were immersed in 98% formic acid for 10 min.

Immunoreactive prions were then detected using the Vectastain avidin-biotin complex (ABC) kit with 3,3′-diaminobenzidine tetrahydrochloride (DAB) as a substrate. Sections were counterstained with haematoxylin before mounting and imaging. Stained sections were imaged using a Brightfield Eclipse Ni-E light microscope (Nikon Instruments Europe BV, Amsterdam, Netherlands) and images captured using Zen 2 software (Carl Zeiss Ltd. Cambridge, United Kingdom), or on a VS120 Olympus slide scanner. For EAE spinal cord sections, whole-slide digital imaging was performed on the VS120 Olympus slide scanner. GFP^+^ was imaged on a confocal microscope (FV3000 Olympus).

### Image Analysis

For the prion disease model, IBA1, GFAP, Luxol Fast Blue (LFB) and disease-specific PrP (PrP^Sc+^) immunostaining was performed and quantified using Fiji/ImageJ software (http://imagej/nih.gov/ij) using the analyse particle algorithm as described previously (Bradford et al., 2022). In the EAE model, DAPI staining was used to identify lesions within white and grey matter infiltrated by immune cells in the spinal cord. GFP^+^, GFAP^+^ and MBP^+^ areas in the spinal cord were quantified using ImageJ (https://imagej.net/). The percentage area of positive staining was calculated using the ‘measure’ tool in ImageJ, following adjustment of the stain specific threshold, which was kept consistent for all mice.

### RNA purification and qRT-PCR analysis

For RNA sequencing, RNA was extracted from brain regions using TRI Reagent according to manufacturer’s instructions. For each extraction ∼100 mg of tissue was processed using 1 ml reagent. RNA pellets were dissolved in UltraPure™ DNAse, RNAse free Distilled Water. RNA quantity was measured using a NanoDrop One. DNase I treatment was used to eliminate genomic DNA contamination of RNA preparations. For qRT-PCR analysis in the prion model, RNA isolation and expression quantification was performed as described previously (Bradford et al., 2022). The primer sequences used are shown in Table 1.

### RNA-seq analysis

Library preparation and sequencing was performed at the University of Queensland Sequencing Facility (University of Queensland, Brisbane, Australia). Bar-coded mRNA-Seq libraries were generated using the Illumina Stranded mRNA Library Prep Ligation kit. The RNA libraries were pooled in equimolar ratios prior to sequencing using the Illumina NovaSeq 6000. Paired-end 102 bp reads were generated using either a NovaSeq 6000 S1 reagent kit v1.5 (200 cycles) or an SP reagent kit v1.5 (200 cycles). After sequencing, fastq files were created using bcl2fastq2 (v2.20.0.422). The sequencing depth ranged from 15-20 million reads per sample. Raw sequencing data (in the form of .fastq files) was provided by the sequencing facility for further analysis.

Raw reads were pre-processed raw reads using fastp v0.23.2 (Chen et al., 2018) and quality controlled using FastQC as described previously (Keshvari et al., 2021).

Transcript expression levels were quantified as transcripts per million using the pseudo-aligner, kallisto (v0.46.0) (Bray et al., 2016). The reference transcriptome used was created by combining the unique protein-coding transcripts from the Ensembl and NCBI RefSeq databases of the GRCm39 annotation, as previously described (Summers et al., 2020). Kallisto expression files were imported into RStudio (R version 4.2.1) using the tximport package (v1.24.0) to collates the transcript-level expression generated by Kallisto into gene-level expression for use by downstream tools.

### Infection with prions and clinical disease assessment

Prion infection was established directly in the CNS by intracerebral (IC) injection into the right medial temporal lobe with 20 µL of a 1% (weight/volume) brain homogenate prepared from mice terminally infected with mouse-passaged ME7 scrapie prions. The mice were coded and assessed blindly at daily intervals for development of clinical signs of prion disease and culled at the times indicated or at an established humane end-point upon the development of terminal clinical signs of prion disease (Donaldson et al., 2020). At the end of the experiment each brain was removed and halved sagittally across the midline. One brain half was immediately flash frozen at the temperature of liquid nitrogen for gene expression or protein analysis, the other brain half was fixed in 10% neutral buffered formalin for histopathology. Brain sections were prepared and haematoxylin & eosin (H&E) stained, and clinical prion disease was confirmed by histopathological assessment of the magnitude and distribution of the spongiform vacuolar degeneration in nine grey matter areas using a 0-5 scale and three white matter areas using a 0-3 scale: G1, dorsal medulla; G2, cerebellar cortex; G3, superior colliculus; G4, hypothalamus; G5, thalamus; G6, hippocampus; G7, septum; G8, retrosplenial and adjacent motor cortex; G9, cingulate and adjacent motor cortex; W1, inferior and middle cerebellar peduncles; W2, decussation of superior cerebellar peduncles; and W3, cerebellar peduncles (Fraser and Dickinson, 1967).

### Western blot detection of prion protein

Prion-specific PrP^Sc^ was detected in flash-frozen brain samples by Western blotting as described previously (Bradford et al., 2017). Briefly, brain homogenates (10% weight/volume) were prepared in NP40 lysis buffer (1% NP40, 0.5% sodium deoxycholate, 150 mM NaCl, 50 mM TrisHCl [pH 7.5]). To detect relatively proteinase-resistant PrP^Sc^ a sample of the homogenate was treated with 20 μg/ml proteinase K (PK) at 37°C for 1 h. The digestion was stopped by addition of 1 mM phenylmethylsulfonyl fluoride. Samples were then denatured by incubation at 85°C for 15 min in 1X SDS sample buffer and separated via electrophoresis using Nupage 12% Tris-glycine polyacrylamide gels. Proteins were then transferred to polyvinylidene difluoride (PVDF) membranes by semi-dry electroblotting and PrP detected using mouse monoclonal antibody BH1 (McCutcheon et al., 2014) and β-actin detected using mouse monoclonal antibody C4. Membranes were subsequently stained with horseradish peroxidase-conjugated goat anti-species-specific antibody and visualised using chemiluminescence. The relative abundance of PrP in each sample on the Western blots was compared by densitometric analysis using ImageJ software. The PrP abundance in each sample was normalised to β-actin and expressed as the percentage PrP relative to the mean value in the PBS-treated controls.

### EAE Induction and Monitoring

The optimised mouse EAE model was described previously (Khan et al., 2014). Briefly, heat-killed Mycobacterium tuberculosis (HKMT) was dissolved in incomplete Freud’s adjuvant at 10mg/ml. Myelin oligodendrocyte glycoprotein (MOG) was dissolved in sterile 1x PBS to give a final concentration of 2mg/ml. A 1:1 ratio of the premade HKMT/IFA and MOG solutions were mixed using microliter syringes (Hamilton, Franklin, MA). Microemulsion of these solutions was regarded as complete when the solution became a creamy white emulsion. Adult female mice were anaesthetised with isofluorane and injected s.c. with a 50μl of the HKMT MOG microemulsion in the right and left flank, close to the draining lymph nodes (100μl total). An i.p. injection of 0.1ml 2ng/μl of pertussis toxin (PTX) (for a total dose of 200ng) was administered. The 200ng PTX i.p injection was repeated 48 h post-immunisation.

Mice were assessed daily for EAE disease symptoms ranging from limp tail to quadriplegia using a 5-point scale with half point graduations as described (Khan et al., 2014). Body weight was measured and the daily body weight percentage from Day 0 was calculated. Mice were given a facial grimace, coat condition, and hunching score. These scores were totalled. A score of 3 in any category, total score of more than 8, or clinical score of 3.5 (bilateral hindlimb and one forelimb paralysis) indicated euthanasia was appropriate.

### Statistics

Statistical significance between groups was tested using Prism 7.0 software (GraphPad, San Diego, United States). Unpaired student’s t-testings of Mann-Whitney t-testing was used to compare two groups (noted in figure legends). For comparisons of 3 or more, two-way ANOVA with subsequent Sidak’s multiple comparison testing was used. For the PrP^Sc^ analysis, datasets were first compared for normality using the Shapiro-Wilk normality test. Differences between groups were then compared using One-Way ANOVA and post-hoc Tukey multiple comparisons tests. Survival curve data for prion infected mice in each treatment group were compared by Log-rank (Mantel-Cox) test. Data are shown as dot plots of individual animal observations with median values indicated by a horizontal bar. Body weight and brain lesion profile data are presented as mean ± SD. *, *P* < 0.05; **, *P* < 0.01; ***, *P* < 0.001.

## Acknowledgements

The generation of the mice was funded by a grant from the Medical Research Council (MRC) UK grant MR/M019969/1 to DAH. This work was supported by Australian National Health and Medical Research Council (NHMRC) Grant GNT1163981 awarded to DAH and KMS. JS was supported by the Australian Research Training Program Stipend and Tuition Fee Offset. The laboratory receives core support from The Mater Foundation. We thank Dr Liviu Bodea for the *Cx3cr1*^GFP/+^ mice and Dr Carly Cullen for providing the MBP antibody. We acknowledge input and expertise from the Biological Resources facility and the Preclinical Imaging, Microscopy, Histology and Flow Cytometry facilities of the Translational Research Institute (TRI) and the Queen’s Medical Research Institute. TRI is supported by the Australian Government. **Figure 6A** was created with biorender.com.

NAM was supported by project (BB/S005471/1) and Institute Strategic Programme Grant funding from the Biotechnology and Biological Sciences Research Council (grant numbers BBS/E/D/10002071, BBS/E/D/20002173 & BBS/E/RL/230002B). RP was supported by a PhD studentship from the Royal (Dick) School of Veterinary Studies (University of Edinburgh, UK).

## Author contributions

JS Investigation, formal analysis, writing

RP Investigation

BMB Investigation, visualization, formal analysis

DCC Formal analysis, data curation

IT Investigation

CP Resources, methodology

NM, TW Resources, methodology, supervision

KMI, KMS Conceptualisation, funding acquisition, supervision, review and editing

NAM Supervision, funding acquisition, review and editing

DAH Investigation, conceptualization, funding acquisition, original draft.

---

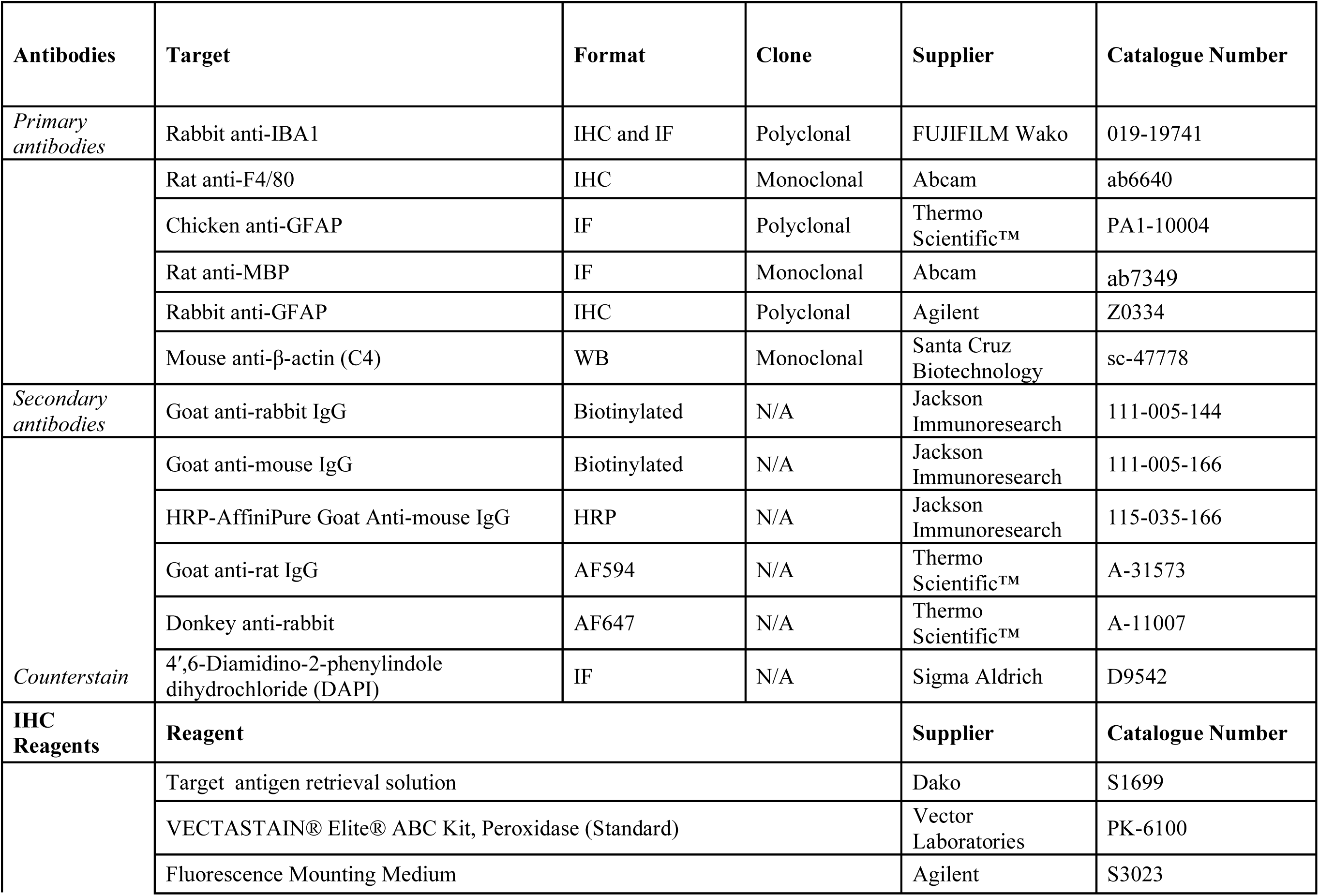

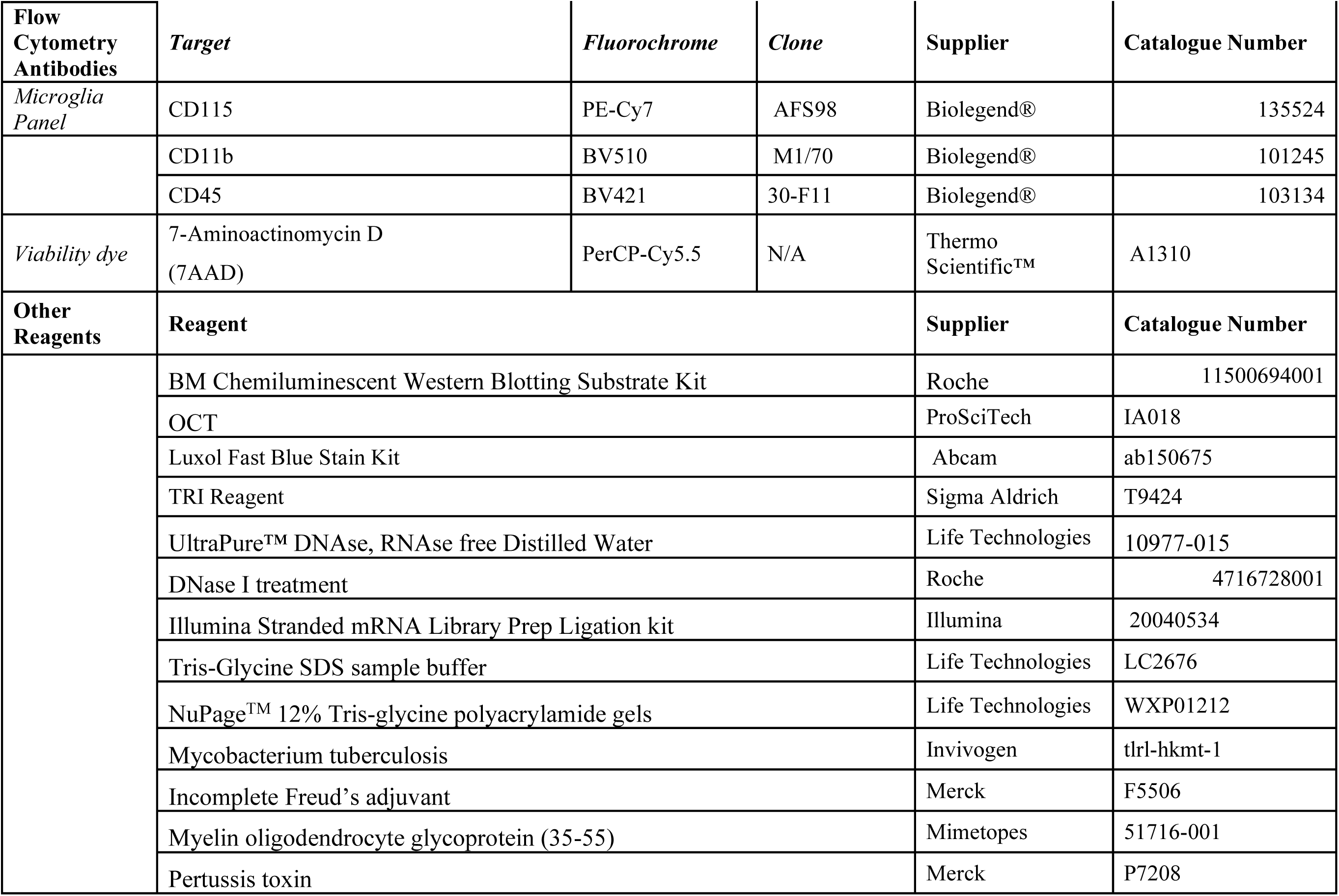

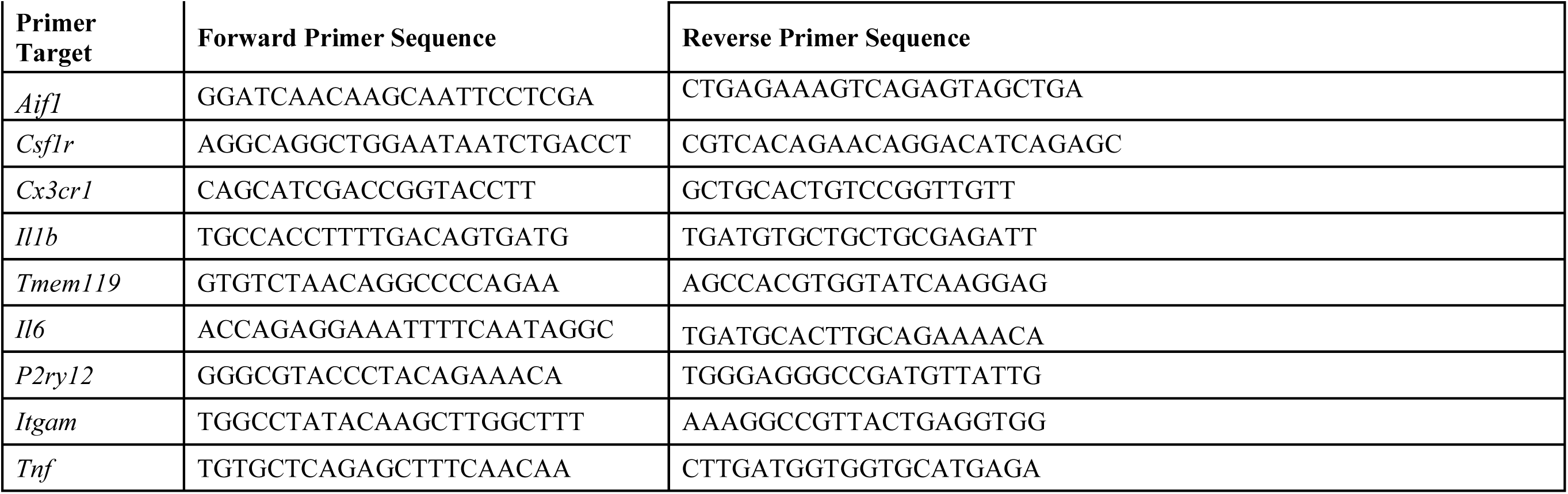

## References

1. Absinta, M., et al., 2021. A lymphocyte-microglia-astrocyte axis in chronic active multiple sclerosis. Nature. 597, 709–714.

2. Aguzzi, A., Zhu, C., 2017. Microglia in prion diseases. J Clin Invest. 127, 3230–3239.

3. Ajami, B., et al., 2011. Infiltrating monocytes trigger EAE progression, but do not contribute to the resident microglia pool. Nat Neurosci. 14, 1142–9.

4. Arreola, M. A., et al., 2021. Microglial dyshomeostasis drives perineuronal net and synaptic loss in a CSF1R(+/-) mouse model of ALSP, which can be rescued via CSF1R inhibitors. Sci Adv. 7, eabg1601.

5. Artegiani, B., et al., 2017. A Single-Cell RNA Sequencing Study Reveals Cellular and Molecular Dynamics of the Hippocampal Neurogenic Niche. Cell Rep. 21, 3271–3284.

6. Askew, K., et al., 2017. Coupled Proliferation and Apoptosis Maintain the Rapid Turnover of Microglia in the Adult Brain. Cell Rep. 18, 391–405.

7. Barki, M., Xue, H., 2022. GABRB2, a key player in neuropsychiatric disorders and beyond. Gene. 809, 146021.

8. Basilico, B., et al., 2022. Microglia control glutamatergic synapses in the adult mouse hippocampus. Glia. 70, 173–195.

9. Basilico, B., et al., 2019. Microglia shape presynaptic properties at developing glutamatergic synapses. Glia. 67, 53–67.

10. Berdowski, W. M., et al., 2022. Dominant-acting CSF1R variants cause microglial depletion and altered astrocytic phenotype in zebrafish and adult-onset leukodystrophy. Acta Neuropathol. 144, 211–239.

11. Bertot, C., et al., 2019. Role of CX3CR1 Signaling on the Maturation of GABAergic Transmission and Neuronal Network Activity in the Neonate Hippocampus. Neuroscience. 406, 186–201.

12. Bittner, S., et al., 2014. Myelin oligodendrocyte glycoprotein (MOG35-55) induced experimental autoimmune encephalomyelitis (EAE) in C57BL/6 mice. J Vis Exp. 51275.

13. Biundo, F., et al., 2023a. Trem2 Enhances Demyelination in the Csf1r(+/-) Mouse Model of Leukoencephalopathy. Biomedicines. 11, 2094.

14. Biundo, F., et al., 2020. Microglial reduction of colony stimulating factor-1 receptor expression is sufficient to confer adult onset leukodystrophy. Glia. 69, 779–791.

15. Biundo, F., et al., 2023b. Elevated granulocyte colony stimulating factor (CSF) causes cerebellar deficits and anxiety in a model of CSF-1 receptor related leukodystrophy. Glia. 71, 775–794.

16. Bourel, J., et al., 2021. Complement C3 mediates early hippocampal neurodegeneration and memory impairment in experimental multiple sclerosis. Neurobiol Dis. 160, 105533.

17. Bradford, B. M., et al., 2022. Microglia deficiency accelerates prion disease but does not enhance prion accumulation in the brain. Glia. 70, 2169–2187.

18. Bradford, B. M., et al., 2017. Oral Prion Disease Pathogenesis Is Impeded in the Specific Absence of CXCR5-Expressing Dendritic Cells. J Virol. 91, e00124–17.

19. Bray, N. L., et al., 2016. Near-optimal probabilistic RNA-seq quantification. Nat Biotechnol. 34, 525–7.

20. Brown, F. N., et al., 2023. Early postnatal microglial ablation in the Ccdc39 mouse model reveals adverse effects on brain development and in neonatal hydrocephalus. Fluids Barriers CNS. 20, 42.

21. Carroll, J. A., et al., 2018. Microglia Are Critical in Host Defense against Prion Disease. J Virol. 92, e00549–18.

22. Chadarevian, J. P., et al., 2024. Therapeutic potential of human microglial transplantation in a chimeric model of CSF1R-related leukoencephalopathy. Neuron. In press.

23. Chen, M. B., et al., 2020. Brain Endothelial Cells Are Exquisite Sensors of Age-Related Circulatory Cues. Cell Rep. 30, 4418–4432 e4.

24. Chen, S., et al., 2018. fastp: an ultra-fast all-in-one FASTQ preprocessor. Bioinformatics. 34, i884–i890.

25. Chitu, V., et al., 2020. Microglial Homeostasis Requires Balanced CSF-1/CSF-2 Receptor Signaling. Cell Rep. 30, 3004–3019 e5.

26. Chitu, V., et al., 2015. Phenotypic characterization of a Csf1r haploinsufficient mouse model of adult-onset leukodystrophy with axonal spheroids and pigmented glia (ALSP). Neurobiol Dis. 74, 219–28.

27. Chitu, V., et al., 2021. Modeling CSF-1 receptor deficiency diseases - how close are we? FEBS J. 289, 5049–5073.

28. Chitu, V., Stanley, E. R., 2017. Regulation of Embryonic and Postnatal Development by the CSF-1 Receptor. Curr Top Dev Biol. 123, 229–275.

29. Cho, H., et al., 2023. Adnp-mutant mice with cognitive inflexibility, CaMKIIalpha hyperactivity, and synaptic plasticity deficits. Mol Psychiatry. 28, 3548–3562.

30. Corsi, G., et al., 2022. Microglia modulate hippocampal synaptic transmission and sleep duration along the light/dark cycle. Glia. 70, 89–105.

31. Cserep, C., et al., 2022. Microglial control of neuronal development via somatic purinergic junctions. Cell Rep. 40, 111369.

32. Dai, X. M., et al., 2002. Targeted disruption of the mouse colony-stimulating factor 1 receptor gene results in osteopetrosis, mononuclear phagocyte deficiency, increased primitive progenitor cell frequencies, and reproductive defects. Blood. 99, 111–20.

33. Diebold, M., et al., 2023. How myeloid cells shape experimental autoimmune encephalomyelitis: At the crossroads of outside-in immunity. Eur J Immunol. 53, e2250234.

34. Donaldson, D. S., et al., 2020. Accelerated onset of CNS prion disease in mice co-infected with a gastrointestinal helminth pathogen during the preclinical phase. Sci Rep. 10, 4554.

35. Elmore, M. R., et al., 2014. Colony-stimulating factor 1 receptor signaling is necessary for microglia viability, unmasking a microglia progenitor cell in the adult brain. Neuron. 82, 380–97.

36. Faust, T. E., et al., 2023. A comparative analysis of microglial inducible Cre lines. Cell Rep. 42, 113031.

37. Forrest, A. R., et al., 2014. A promoter-level mammalian expression atlas. Nature. 507, 462–70.

38. Fournier, A. P., et al., 2023. Single-Cell Transcriptomics Identifies Brain Endothelium Inflammatory Networks in Experimental Autoimmune Encephalomyelitis. Neurol Neuroimmunol Neuroinflamm. 10, e200046.

39. Fraser, H., Dickinson, A. G., 1967. Distribution of experimentally induced scrapie lesions in the brain. Nature. 216, 1310–1.

40. Freeman, T. C., et al., 2022. Graphia: A platform for the graph-based visualisation and analysis of high dimensional data. PLoS Comput Biol. 18, e1010310.

41. Gomez-Nicola, D., et al., 2013. Regulation of microglial proliferation during chronic neurodegeneration. J Neurosci. 33, 2481–93.

42. Green, K. N., et al., 2020. To Kill a Microglia: A Case for CSF1R Inhibitors. Trends Immunol. 41, 771–784.

43. Guneykaya, D., et al., 2018. Transcriptional and Translational Differences of Microglia from Male and Female Brains. Cell Rep. 24, 2773–2783 e6.

44. Guo, L., Ikegawa, S., 2021. From HDLS to BANDDOS: fast-expanding phenotypic spectrum of disorders caused by mutations in CSF1R. J Hum Genet. 66, 1139–1144.

45. Gyoneva, S., et al., 2019. Cx3cr1-deficient microglia exhibit a premature aging transcriptome. Life Sci Alliance. 2, e201900453.

46. Hagan, N., et al., 2020. CSF1R signaling is a regulator of pathogenesis in progressive MS. Cell Death Dis. 11, 904.

47. Hanamsagar, R., et al., 2018. Generation of a microglial developmental index in mice and in humans reveals a sex difference in maturation and immune reactivity. Glia. 66, 460.

48. Haney, M. S., et al., 2024. APOE4/4 is linked to damaging lipid droplets in Alzheimer’s disease microglia. Nature. 628, 154–161.

49. Hume, D. A., 2023. Fate-mapping studies in inbred mice: A model for understanding macrophage development and homeostasis? Eur J Immunol. 53, e2250242.

50. Hume, D. A., et al., 2020. Phenotypic impacts of CSF1R deficiencies in humans and model organisms. J Leukoc Biol. 107, 205–219.

51. Hume, D. A., et al., 2023. Macrophage biology in the single cell era: facts and artefacts. Blood. 142, 1339–1347.

52. Hwang, D., et al., 2022. CSF-1 maintains pathogenic but not homeostatic myeloid cells in the central nervous system during autoimmune neuroinflammation. Proc Natl Acad Sci U S A. 119, e2111804119.

53. Jacquelot, N., et al., 2022. Innate lymphoid cells and cancer. Nat Immunol. 23, 371–379.

54. Jung, S., et al., 2000. Analysis of fractalkine receptor CX(3)CR1 function by targeted deletion and green fluorescent protein reporter gene insertion. Mol Cell Biol. 20, 4106–14.

55. Kempthorne, L., et al., 2020. Loss of homeostatic microglial phenotype in CSF1R-related Leukoencephalopathy. Acta Neuropathol Commun. 8, 72.

56. Keshvari, S., et al., 2021. CSF1R-dependent macrophages control postnatal somatic growth and organ maturation. PLoS Genet. 17, e1009605.

57. Khan, N., et al., 2014. Establishment and characterization of an optimized mouse model of multiple sclerosis-induced neuropathic pain using behavioral, pharmacologic, histologic and immunohistochemical methods. Pharmacol Biochem Behav. 126, 13–27.

58. Kiani Shabestari, S., et al., 2022. Absence of microglia promotes diverse pathologies and early lethality in Alzheimer’s disease mice. Cell Rep. 39, 110961.

59. Kim, J. D., et al., 2019. Microglial UCP2 Mediates Inflammation and Obesity Induced by High-Fat Feeding. Cell Metab. 30, 952–962 e5.

60. Konno, T., et al., 2018. CSF1R-related leukoencephalopathy: A major player in primary microgliopathies. Neurology. 91, 1092–1104.

61. Konno, T., et al., 2017. Clinical and genetic characterization of adult-onset leukoencephalopathy with axonal spheroids and pigmented glia associated with CSF1R mutation. Eur J Neurol. 24, 37–45.

62. Lauro, C., et al., 2008. Activity of adenosine receptors type 1 Is required for CX3CL1-mediated neuroprotection and neuromodulation in hippocampal neurons. J Immunol. 180, 7590–6.

63. Lelios, I., et al., 2020. Emerging roles of IL-34 in health and disease. J Exp Med. 217, e20190290.

64. Liu, Y. J., et al., 2021. Microglia Elimination Increases Neural Circuit Connectivity and Activity in Adult Mouse Cortex. J Neurosci. 41, 1274–1287.

65. McCutcheon, S., et al., 2014. Prion protein-specific antibodies that detect multiple TSE agents with high sensitivity. PLoS One. 9, e91143.

66. McLay, R. N., et al., 1997. Granulocyte-macrophage colony-stimulating factor crosses the blood--brain and blood--spinal cord barriers. Brain. 120 (Pt 11), 2083–91.

67. McNamara, N. B., et al., 2023. Microglia regulate central nervous system myelin growth and integrity. Nature. 613, 120–129.

68. Montilla, A., et al., 2023. Microglia and meningeal macrophages depletion delays the onset of experimental autoimmune encephalomyelitis. Cell Death Dis. 14, 16.

69. Munji, R. N., et al., 2019. Profiling the mouse brain endothelial transcriptome in health and disease models reveals a core blood-brain barrier dysfunction module. Nat Neurosci. 22, 1892–1902.

70. Munro, D. A. D., et al., 2024. Microglia provide resilience against region-selective neuropathology with ageing. Neuron. In press.

71. Nissen, J. C., et al., 2018. Csf1R inhibition attenuates experimental autoimmune encephalomyelitis and promotes recovery. Exp Neurol. 307, 24–36.

72. Obst, J., et al., 2017. The Role of Microglia in Prion Diseases: A Paradigm of Functional Diversity. Front Aging Neurosci. 9, 207.

73. Oiso, N., et al., 2013. Piebaldism. J Dermatol. 40, 330–5.

74. Oosterhof, N., et al., 2019. Homozygous Mutations in CSF1R Cause a Pediatric-Onset Leukoencephalopathy and Can Result in Congenital Absence of Microglia. Am J Hum Genet. 104, 936–947.

75. Oosterhof, N., et al., 2018. Colony-Stimulating Factor 1 Receptor (CSF1R) Regulates Microglia Density and Distribution, but Not Microglia Differentiation In Vivo. Cell Rep. 24, 1203–1217 e6.

76. Pan, L., et al., 2023. Oligodendrocyte-lineage cell exocytosis and L-type prostaglandin D synthase promote oligodendrocyte development and myelination. Elife. 12, e77441.

77. Paolicelli, R. C., et al., 2022. Microglia states and nomenclature: A field at its crossroads. Neuron. 110, 3458–3483.

78. Patkar, O. L., et al., 2021. Analysis of homozygous and heterozygous Csf1r knockout in the rat as a model for understanding microglial function in brain development and the impacts of human CSF1R mutations. Neurobiol Dis. 151, 105268.

79. Pridans, C., et al., 2013. CSF1R mutations in hereditary diffuse leukoencephalopathy with spheroids are loss of function. Sci Rep. 3, 3013.

80. Race, B., et al., 2022. Microglia have limited influence on early prion pathogenesis, clearance, or replication. PLoS One. 17, e0276850.

81. Rademakers, R., et al., 2011. Mutations in the colony stimulating factor 1 receptor (CSF1R) gene cause hereditary diffuse leukoencephalopathy with spheroids. Nat Genet. 44, 200–5.

82. Reith, A. D., et al., 1990. W mutant mice with mild or severe developmental defects contain distinct point mutations in the kinase domain of the c-kit receptor. Genes Dev. 4, 390–400.

83. Rogers, J. T., et al., 2011. CX3CR1 deficiency leads to impairment of hippocampal cognitive function and synaptic plasticity. J Neurosci. 31, 16241–50.

84. Rojo, R., et al., 2019. Deletion of a Csf1r enhancer selectively impacts CSF1R expression and development of tissue macrophage populations. Nat Commun. 10, 3215.

85. Sasmono, R. T., et al., 2003. A macrophage colony-stimulating factor receptor-green fluorescent protein transgene is expressed throughout the mononuclear phagocyte system of the mouse. Blood. 101, 1155–63.

86. Stables, J., et al., 2022. A kinase-dead Csf1r mutation associated with adult-onset leukoencephalopathy has a dominant inhibitory impact on CSF1R signalling. Development. 149, dev200237.

87. Stanley, E. R., Chitu, V., 2014. CSF-1 receptor signaling in myeloid cells. Cold Spring Harb Perspect Biol. 6, a021857.

88. Summers, K. M., et al., 2020. Transcriptional network analysis of transcriptomic diversity in resident tissue macrophages and dendritic cells in the mouse mononuclear phagocyte system. PLoS Biol. 03.24., e3000859.

89. Tada, M., et al., 2016. Characteristic microglial features in patients with hereditary diffuse leukoencephalopathy with spheroids. Ann Neurol. 80, 554–65.

90. Villa, A., et al., 2018. Sex-Specific Features of Microglia from Adult Mice. Cell Rep. 23, 3501–3511.

91. Wiedrick, J., et al., 2021. Sex differences in EAE reveal common and distinct cellular and molecular components. Cell Immunol. 359, 104242.

92. Yousef, H., et al., 2019. Aged blood impairs hippocampal neural precursor activity and activates microglia via brain endothelial cell VCAM1. Nat Med. 25, 988–1000.

93. Yue, X., et al., 1993. Expression of mRNA encoding the macrophage colony-stimulating factor receptor (c-fms) is controlled by a constitutive promoter and tissue-specific transcription elongation. Mol Cell Biol. 13, 3191–201.

